# Recurrent training rejuvenates and enhances transcriptome and methylome responses in young and older human muscle

**DOI:** 10.1101/2020.06.30.179465

**Authors:** Sara Blocquiaux, Monique Ramaekers, Ruud Van Thienen, Henri Nielens, Christophe Delecluse, Katrien De Bock, Martine Thomis

**Affiliations:** Physical Activity, Sport & Health research group, Department of Movement Sciences, KU Leuven, Belgium; Exercise Physiology research group, Department of Movement Sciences, KU Leuven, Belgium; Research group for Neurorehabilitation, Department of Rehabilitation Sciences, KU Leuven, Belgium; Saint-Luc University Hospital and Institute of NeuroScience, System and Cognition Division, UCLouvain, Belgium; Laboratory of Exercise and Health, Department of Health Sciences and Technology, ETH Zürich, Switzerland

**Keywords:** DNA methylation, RNA expression, muscle ageing, immobilization, detraining, epi-memory

## Abstract

**Background:** The interaction between the muscle methylome and transcriptome is understudied during ageing and periods of resistance training in young, but especially older adults. In addition, more information is needed on the role of retained methylome training adaptations in muscle memory to understand muscle phenotypical and molecular restoration after inactivity or disuse.

**Methods:** We measured CpG methylation (microarray) and RNA expression (RNA sequencing) in young (n = 5; age = 22 ± 2 yrs) and older (n = 6; age = 65 ± 5 yrs) vastus lateralis muscle samples, taken at baseline, after 12 weeks of resistance training, after training interruption (2 weeks of leg immobilization in young men, 12 weeks of detraining in older men) and after 12 weeks of retraining to identify muscle memory-related adaptations and rejuvenating effects of training.

**Results:** We report that of the 427 differentially expressed genes with advanced age, 71 % contained differentially methylated (dm)CpGs in older versus young muscle. The more dmCpGs within a gene, the clearer the inverse methylation-expression relationship. Around 73 % of the age-related dmCpGs approached younger methylation levels when older muscle was trained for 12 weeks. A second resistance training period after training cessation increased the number of hypomethylated CpGs and upregulated genes in both young and older muscle. We found indication for an epi-memory within pro-proliferating *AMOTL1* in young muscle and mechanosensing-related *VCL* in older muscle. For the first time, we integrate muscle methylome and transcriptome data in relation to both ageing and training/inactivity-induced responses and identify focal adhesion as an important pathway herein.

**Conclusion:** Previously trained muscle is more responsive to training than untrained muscle at methylome and transcriptome level and recurrent resistance training can partially restore ageing-induced methylome alterations.

## INTRODUCTION

Age-related skeletal muscle decay is the consequence of primary and secondary ageing.^1^ The first is described as the inevitable deterioration of cellular structures and biological functions with little chance of reversal and inter-individual differences caused by genetic factors.^2^ Secondary ageing, on the other hand, is the result of harmful lifestyle choices and environmental factors. These harmful factors can influence epigenetic mechanisms, defined as events that regulate gene expression without changing the DNA sequence. One of the most studied epigenetic events is DNA methylation^3^, which mostly refers to CpG methylation and implies the binding of a methyl group to a 5’-cytosin-phosphate-guanine-3’ dinucleotide (CpG). Hypermethylation of CpG-dense promoter regions of actively transcribed genes has been associated with transcriptional repression, also known as ‘gene silencing’.^4^ The methylome of ageing tissues is characterized by global CpG hypomethylation, loci-specific differential methylation and more divergence between DNA methylation patterns of different individuals with the end result leading to aberrant gene expression, reactivation of transposable elements and genomic instability.^5,6^ Age-related methylation changes in muscle tissue, on the other hand, appear distinct compared to other tissues^7^. Recently, two genome-wide DNA methylation analyses identified 5,518^8^ and 17,994^9^ differentially methylated CpGs (dmCpGs) between old and young skeletal muscle with most of them hypermethylated in older adults. *The integration between age-related muscle methylome and transcriptome changes remains yet to be studied and is a crucial step in understanding the molecular mechanism behind muscle ageing*.

As DNA methylation modifications are reversible, adopting a healthy lifestyle could potentially restore the age-associated damage to the muscle methylome and promoting physical activity is one strategy to do this. For instance, endurance exercise interventions are associated with altered methylation levels in genes involved in metabolism, cardiovascular adaptations, inflammatory and immunological processes and muscle remodelling in young and middle-aged muscle.^10–13^ These beneficial methylome adaptations could even be passed on to later generations, as has been observed in mice^14^. In older muscle, lifelong physical activity induced promoter hypomethylation of genes involved in energy metabolism, myogenesis, contractile properties and oxidative stress.^15^ The impact of resistance training (RT) on the muscle methylome is less documented in literature, especially in elderly, but it has been suggested that RT has a distinct effect on the methylation status of, among other, growth-related genes^13,16^. *Understanding the impact of RT on the muscle methylome is important, as resistance exercises can contribute to regaining muscle mass and strength and should be part of the exercise program designed to counteract muscle ageing*.

Nevertheless, lifelong training without training interruptions seems utopian as training can be occasionally ceased for various reasons. Fortunately, training-related muscle adaptations, and especially strength gains, are preserved longer than the initial training period and are often more rapidly regained following retraining.^17–19^ In addition, the basal and acute RT-induced transcription of key genes involved in muscular adaptations appear to be influenced by a previous RT period.^20^ This so-called muscle memory has been partially attributed to motor learning^18^ and linked to the retention of myonuclei during muscle atrophy^21^, as well as to lasting epigenetic modifications^16^. The slow reversibility of methylation has been, for instance, seen when methylation changes in young skeletal muscle following five days of high-fat diet only partially and non-significantly reversed after six to eight weeks^22^. The epigenetic retention of prior external stimuli in a way that re-encountering these stimuli later in life is associated with an enhanced muscle phenotypical response compared to the initial response (i.e. *epi-memory*^23^), has been described in relation to metabolic (e.g. obesity^24^), catabolic (e.g. TNF-α^25^) and anabolic (e.g. RT^16^) stimuli. As DNA methylation modifications are transient in nature after one bout of exercise^26^, the question remains how long beneficial methylation alterations due to an extended period of RT are retained and whether multiple re-encounters are necessary to accumulate and stabilize these changes. Recently, a number of genes were identified in young skeletal muscle with a possible ‘epi-memory’.^16,27^ These genes displayed hypomethylation after seven weeks of RT and continued to be hypomethylated during seven weeks of detraining, leading to enhanced gene expression following retraining. The number of hypomethylated dmCpGs following retraining was also twice the number compared to following training, which coincided with larger muscle growth.^16^ *Our knowledge on epi-memory is, however, still rudimentary and identifying genes that constitute the epi-memory might provide crucial insights for the future development of therapies that promote muscle regain after inactivity*.

Here, we use a training, immobilization and retraining resistance exercise protocol in young men to identify muscle memory-related genes at muscle methylome and transcriptome level. In addition, we test the rejuvenating effect of RT on the older skeletal muscle methylome and whether these beneficial methylome adaptations are maintained during detraining and promote an enhanced transcriptional response during retraining.

## METHODS

### Subjects and experimental design

Nine older men, divided in an exercise group (n = 6, age: 66 ± 5 years, BMI: 26 ± 3 kg/m^2^; mean ± SD) and control group (n = 3, age: 70 ± 4 years, BMI: 26 ± 2 kg/m^2^), and five younger men (age: 22 ± 2 years, BMI: 24 ± 3 kg/m^2^) gave written consent for this study after being fully informed. All men were non-smoking, untrained, healthy individuals (exclusion factors described in detail elsewhere^28^). The experimental design of the study is displayed in **Figure 1**. One younger subject withdrew after the first training period for reasons unrelated to this investigation. Baseline and post-training samples from this subject are included in the analysis. The study protocol was approved by the Medical Ethics Committee of the Catholic University of Leuven (S59380) and is in accordance with the Declaration of Helsinki.

**Figure 1.**
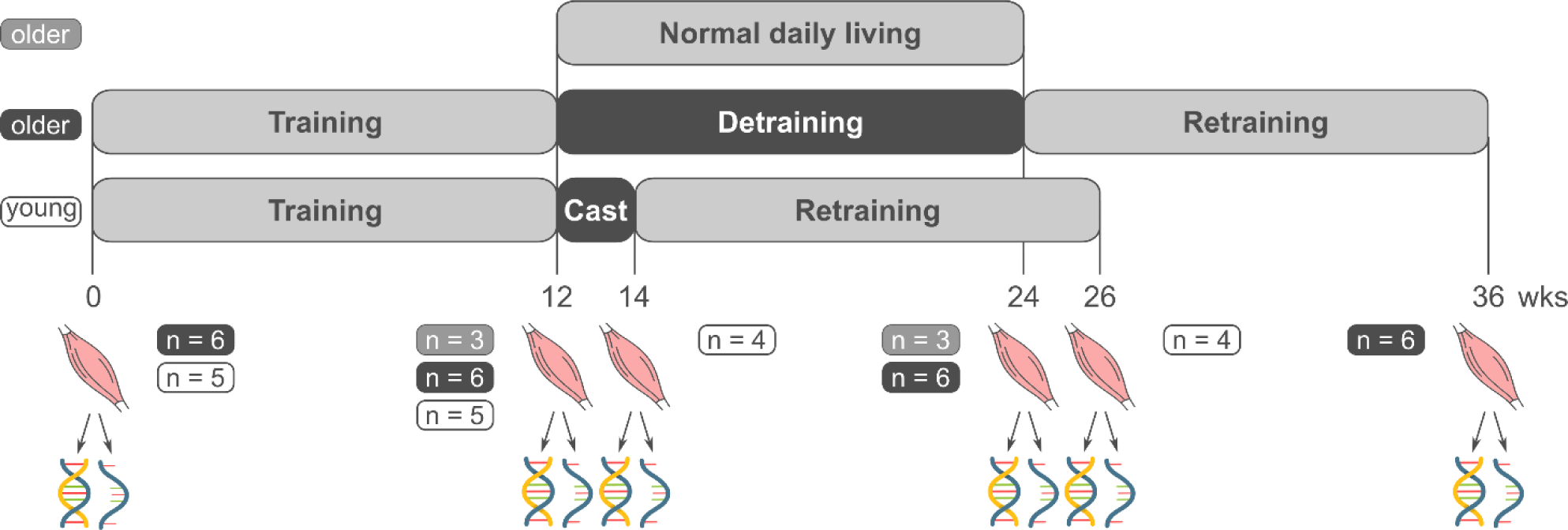
Experimental design. Five younger and six older subjects participated in a 12-week RT program (training). Thereafter, the right leg of the younger subjects was immobilized for 2 weeks (cast), whereas the older subjects ceased training for 12 weeks (detraining). A second 12-week RT-period was provided following the casting/detraining phase (retraining). Muscle biopsies (image of a muscle) were taken before training, after training, after casting/detraining and after retraining. Three older subjects provided a muscle sample from before and after a 12-week period of normal daily living (control group). Part of the biopsy was used to extract DNA and measure DNA methylation (image of a double strand). The other part was used to extract RNA and perform RNA expression analysis (image of a single strand).

### Resistance training program

A 12-week supervised RT-program was conducted as previously described^28^. Briefly, three times a week, on non-consecutive days, subjects performed eight exercises targeting all major muscle groups of the upper and lower body with load intensity gradually increasing every four weeks (65 % to 80 % of 1-repetition maximum). The training load was increased for each exercise when the final set comprised more repetitions than the prescribed number. The vastus lateralis muscle was specifically trained with a Signature Series Plate Loaded Linear Leg Press and Insignia Series Leg Extension machine (Life Fitness, Barendrecht, the Netherlands). The same RT program was used for training and retraining phases with starting loads adapted to the physical state at that time.

### Training interruption

#### Young subjects

The right leg of the young subjects was immobilized with the knee in nearly full extension for two weeks using a full leg cast (foot included). Subjects were instructed to walk with crutches during the immobilization phase. To prevent deep vein thrombosis, subjects were screened for risk factors and anticoagulant enoxaparin (40 mg per 0.4 ml per day, Clexane®, Sanofi Belgium, Diegem, Belgium) was administered daily via subcutaneous injection into the abdominal skinfold.

#### Older subjects in the exercise group

The training program was interrupted by a detraining phase. Subjects were instructed not to engage in any structured strength or endurance exercises for 12 weeks and to resume normal physical activities from before the start of the study. To verify this, subjects weekly filled in a physical activity diary.

### Skeletal muscle biopsy

Muscle biopsies were collected at baseline (left leg), ∼72 h after the final RT session of the first training period (left leg), at the end of week 12 from the detraining period (older subjects, right leg) or immediately after cast removal (younger subjects, right leg) and ∼72 h after the final RT session of the retraining period (right leg). The older control subjects provided samples at the post-training (right leg) and post-detraining (left leg) time point of the exercise group. Samples were taken under local anaesthesia from the mid-portion of the m. vastus lateralis with a 5-mm Bergström needle^29^, immediately cleaned of fat and connective tissue and frozen in liquid nitrogen before being stored at -80°C for further analysis.^28^ Subjects were asked not to participate in moderate to vigorous physical activity and not to consume alcohol in the 24 h before biopsy collection. Food intake during the 72 h before biopsy collection was comparable between each time point within each subject.

### DNA extraction and Illumina methylation assay

Genomic DNA was extracted from 15 to 20 mg muscle tissue with the proteinase K method, which uses proteinase K and lysis buffer. DNA was quantitated with the Qubit 2.0 Fluorometer (Invitrogen). DNA samples were transferred to the Genomics Core facility (UZ/KULeuven, Belgium) to measure DNA methylation level at over 850,000 CpGs using the Illumina Infinium MethylationEPIC BeadChip.

### Pre-processing DNA methylation data

IDAT-files, containing raw intensity data from the EPIC beadchip, were pre-processed in R (version 3.6.1)^30^. Firstly, the R/Bioconductor package *minfi* v1.30.0^31^ was used to read the IDAT-files and perform quality control. None of the samples contained more than 10 % of sites with detection p-value > 0.05^32^ and all samples were considered good quality samples based upon clustering the log median intensities of raw methylated and unmethylated values. Background correction was performed with Noob dye-bias normalization^33^, implemented in *minfi*. Next, β-values, defined as the ratio of the methylated probe intensity and the overall intensity, with β = 0 indicating sites are completely unmethylated and β = 1 expressing fully methylated sites^34^, were extracted from the normalized data. These β-values were subsequently normalized with BMIQ^35^, implemented in R/Bioconductor package *wateRmelon* v1.28.0^36^, to correct for the different design of type I and type II probes in the EPIC beadchip, as suggested by Liu et al.^37^. Finally, we filtered out 357 probes that had failed in 20 % of the samples based on detection p-value > 0.05^32^, 20,655 probes that had failed to hybridize based on a bead count < 3 in 5 % of the samples^32^, 97,253 probes that are cross-reactive or overlap with genetic variants based on the h19 genome^38^ and 206 probes with unknown genomic location. Ultimately, 747,741 probes were considered for downstream analyses, which was performed on M-values, the log2 ratio of the intensities of methylated probe versus unmethylated probe^39^. Principal component analysis (PCA) plots were able to cluster young and older samples based on age, sample ID and timing in the intervention (**Supporting Information Figures S1-2**). To correct for unwanted sources of variation, such as batch effects and cell type heterogeneity within muscle tissue, we used the package *sva* v.3.32.1^40^ to perform surrogate variable analysis (SVA) on the M-values of each subject group, while protecting the information coming from the intervention (variable of interest, included in the full model) and the paired subject IDs (adjustment variable, included in the null and full model). The number of surrogate variables per group was estimated with the default ‘be’ approach^41^. No SVA was performed when comparing young to older muscle methylome, as the function could not handle a design where individuals were nested within groups.

### Differential DNA methylation analysis

Differential DNA methylation analysis was performed in Partek Genomics Suite V.7.0 (Partek Inc., St. Louis, MO, USA). To identify age-related dmCpGs, we applied a mixed model with methylation M-values as response variable, ‘time’ (baseline, post-training) and ‘group’ (younger subjects, older subjects of the exercise group) as fixed factors and ‘subject ID’ as random factor. We specifically interpreted the planned contrasts ‘younger versus older subjects at baseline’ and ‘younger subjects at baseline versus older subjects at post-training’. A CpG list was generated using Benjamini-Hochberg’s^42^ false discovery rate (FDR) < 0.1 and difference in M-values > 0.4^39^ as cut-off rules. To determine within-group differences over time, three separate mixed models were fitted (one for each subject group). Methylation M-values were entered into the model as response variable, ‘time’ (all available time points) and surrogate variables as fixed factors and ‘subject ID’ as random factor. Contrasts were specified to study the different combinations of the various time points. Since only one CpG remained after FDR-control and this part of the study is considered explorative in nature^43^, statistical significance was determined at p < 0.01 with a M-value difference threshold of 0.4. Probes were annotated based on ‘HumanMethylation850’ reference, ‘MethylationEPIC_v-1-0_B4’ annotation file and hg19 genome build. Overlapping genes and gene regions were determined based on Ensembl Transcripts data release 75. “Promoter regions” are defined in this study as the combination of TSS1500 (1500 – 201 base pairs upstream from the transcription start site), TSS200 (200 – 0 base pairs upstream from the transcription start site) and the first exon (5’UTR + the first coding sequence). “Intragenic” will be used as the overlapping term for all exons (except for the first coding sequence), introns, regions between the transcription start and end site of non-coding transcripts and 3’UTR regions. “Intergenic regions” are regions between genes, not including the promoter region. The relationship to CpG islands (island, shore, shelf, open sea) was determined with the UCSC genome browser. We furthermore performed pathway enrichment via a Fisher’s Exact test in Partek Pathway (Partek Inc., St. Louis, MO, USA) based on the KEGG pathway database. We restricted the analysis to pathways of minimal 10 and maximal 500 genes, as small pathways are redundant with larger pathways and larger pathways overly general (e.g. pathways in cancer), inflating their statistical significance^44^. Finally, to find overlapping CpGs, genes and pathways following each intervention phase, we used the Venn diagram tool available via http://bioinformatics.psb.ugent.be/webtools/Venn/. This tool was also consulted to find overlap between dmCpGs of the present study and comparable studies. Pearson correlations tests and Fisher’s exact tests were respectively used to identify correlations and associations between the overlapping datasets (with significance level set at p < 0.05).

### RNA extraction and sequencing

Total RNA was homogenized from 15 to 20 mg of muscle tissue with silicon beads in 1ml Trizol reagent (Invitrogen) using a FastPrep-24 5G (MP Biomedicals) and extracted with phase separation reagent 1-Bromo-3-chloropropane. RNA concentration and purity (A260/230) were determined with SimpliNano (Biochrom). RNA integrity was assessed with the Agilent 2100 Bioanalyzer System and was at least 7.7 in all samples. At the Genomics Core facility, sequencing libraries were prepared with the QuantSeq 3’ mRNA-Seq Library Prep Kit FWD (Lexogen) and sequenced on a HiSeq 4000 Sequencing System (Illumina) using the HiSeq3000/4000 SR Cluster and SBS Kit (Illumina).

### Pre-processing RNA-sequencing data

Quality control of raw reads was performed with FastQC v0.11.5. Adapters were filtered with ea-utils v1.2.2.18. Splice-aware alignment was performed with the CRAN package *Star* v2.6.1b against the human reference genome hg19 based on annotations from Ensembl release 75. The number of allowed mismatches was two. Reads that mapped to more than one site to the reference genome were discarded. The minimal score of alignment quality to be included in count analysis was 10. Resulting SAM and BAM alignment files were handled with Samtools v0.1.19.24^45^. Quantification of reads per gene was performed with HT-Seq count v0.5.3p3^46^. Further pre-processing was performed in R (version 3.6.1)^30^. Data quality assessment and count-based differential expression analysis of the 56,638 sequenced genes were done with R/Bioconductor package *DESeq2* v1.26^47^. By inspecting heatmaps of the count matrix and PCA plots, we detected and excluded one outlier within the older exercise group (post-retraining time point) (**Supporting Information Figure S3**). The *sva* package (svaseq-function) was used in a similar way as described above to detect unwanted variation in the normalized counts. Low count genes were filtered with *DESeq2*’s independent filtering technique, leaving 23,296, 24,661 and 19,211 genes left for expression analysis in the young exercise, older exercise and older control groups, respectively. Results were generated on normalized counts, corrected for library size.

### Differential RNA expression analysis

To test for any difference over the multiple time points, a likelihood ratio test was performed via *DESeq2*, including ‘time’, surrogate variables and ‘subject ID’ in the full model and surrogate variables and ‘subject ID’ in the reduced model. Pairwise comparisons between time points were analysed with Wald tests. Reported p-values were adjusted for multiple testing with the Benjamini-Hochberg procedure^42^ after assigning weights using Independent Hypothesis Weighting (*IHW* package v1.6.0^48^). Differentially expressed genes (DEGs) are reported if FDR < 0.1 and the average read count ≥ 5. Overlap between methylation and expression data was determined with the Venn diagram tool (cfr. supra).

### Comparison with the MetaMEx meta-analysis

We ran the top 100 DEGs of each analysis in MetaMEx^49^, a tool combining transcriptome data from more than 90 human studies that allows to meta-analyse changes in single genes across exercise and disuse studies. The training and immobilization data of the younger men was, respectively, cross-referenced with RT studies of 10 to 20 weeks (n = 58)^50–55^ (**File S4h**) and immobilization/bed rest studies of 5 to 21 days (n = 42)^10,56–59^ (**File S4i**) in healthy, sedentary/active, lean/overweight, younger men. MetaMEx does not include retraining studies. Therefore, we report genes that display the same expression response as RT studies in MetaMEx or the opposite expression response compared to disuse studies in MetaMEx (**File S4j**). We cross-referenced the (re-)training data of the older men with RT studies of 12 to 24 weeks in healthy, sedentary and active, lean or overweight, elderly men (n = 66)^50,52,54,60–62^ (**File S2h, S2j**). MetaMEx does not include detraining studies, instead the detraining data of the older men is compared to a five-day bed rest study in elderly men (n = 10)^56^ (**File S2i**).

## RESULTS

### More hypomethylated CpGs and upregulated genes in previously trained young muscle

A first training period affected methylation levels of 3,623 CpGs in young skeletal muscle (p < 0.01, M-value difference > 0.4) with an approximate 50/50-distribution of hypo-versus hypermethylated CpGs (1,822 vs 1,801) (**Figure 2a, File S4a**). The dmCpGs were not over-or underrepresented in any gene region or structure (**Figure 2b**), thereby mirroring the distribution of the CpGs in the whole genome (**File S4d**). Interestingly, within promoter-associated CpG islands a higher number of CpGs were hypomethylated (1,326) compared to hypermethylated (909), whereas in other areas the number of hypo-versus hypermethylated CpGs was more equally distributed (**Figure 2b**). At transcriptional level, 11 genes were upregulated and 16 genes were downregulated following training (FDR < 0.1, average read count ≥ 5) (**Figure 2a, File S4h**). Ten of these 27 genes were differentially expressed in the same direction as comparable training studies (cfr. supra: Methods) included in the metaMEx meta-analysis (**File S4h**).

**Figure 2.**
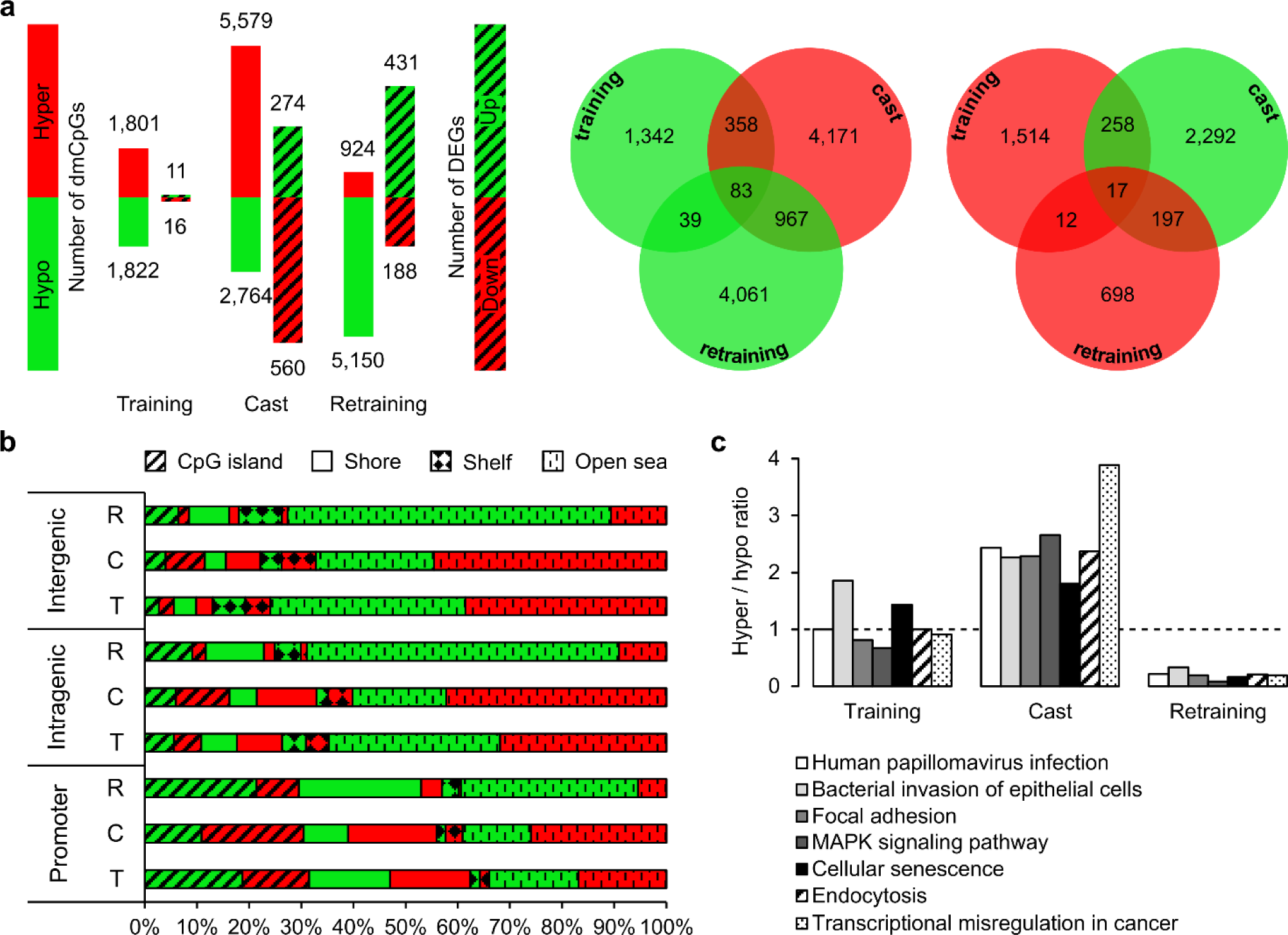
**(a)** Number of hypo- and hypermethylated CpGs (p < 0.01, M-value difference > 0.4; no pattern) versus up- and downregulated genes (DEGs) (FDR < 0.1, average read count ≥ 5; dashed pattern) following training (n = 5), casting (n = 4) and retraining (n = 4) compared to previous time point in young muscle. Venn-diagrams displaying overlap between dmCpGS following training, casting and retraining (hypo- = green; hyper- = red) **(b)** Distribution of dmCpGs (% of the total, hypo- = green; hyper- = red) following training (T), casting (C) and retraining (R) in young muscle across gene regions and gene structures. **(c)** The hyper/hypo ratio of the seven overlapping pathways following training, casting and retraining. This ratio is calculated by dividing the number of hypermethylated CpGs with the number of hypomethylated CpGs. The dashed line (ratio = 1) delimitate an equal level of hyper- and hypomethylated CpGs within a pathway.

The two-week casting period reversed methylation levels in 716 of the 3,623 dmCpGs following training (p < 0.01, M-value difference > 0.4) (**Figure 2a, File S4b**). An additional 7,627 unique CpGs were differentially methylated following the two weeks of immobilization compared to post-training with the majority being hypermethylated (5,138). This immobilization-induced hypermethylation was also reflected within the different gene regions and structures (**Figure 2b**). Two hundred seventy-four genes were upregulated and 560 genes were downregulated following immobilization compared to post-training (FDR < 0.1, average read count ≥ 5) (**Figure 2a, File S4i**). Immobilization has a clear and distinct effect on the muscle transcriptome, as the top 100 most DEGs following immobilization in the present study corresponded largely to what is found in other disuse studies of the MetaMEx meta-analysis (**File S4i**). Most genes within this top 100 were downregulated following immobilization and related to metabolism (49) or muscle structure and contraction (17) (**File S4i**).

A second 12-week training period had a clear hypomethylating effect as 5,150 of the 6,074 dmCpGs displayed lower methylation levels following retraining compared to post-immobilization (p < 0.01, M-value difference > 0.4) (**Figure 2a, File S4c**). Retraining reversed methylation levels in 1,264 of the 8,343 dmCpGs following immobilization (**Figure 2a**). Furthermore, merely 151 CpGs were differentially methylated in the same direction following retraining as following training (**Figure 2a**) and in both CpG-rich and -poor gene areas the frequency of hypomethylated CpGs was enhanced compared to following training (**Figure 2b**). A similar enhanced retraining response was reported earlier in young men after seven weeks detraining^16^. The enhanced retraining response was also reflected at transcriptional level with 431 genes upregulated and 188 downregulated (FDR < 0.1, average read count ≥ 5) (**Figure 2a, File S4j**). Retraining reversed expression levels in 463 of the 834 immobilization-induced transcriptional dysregulated genes. Restoration was observed in, among other, the mitochondrial respiratory chain (16 members of the NADH ubiquinone oxidoreductase complex, 8 members of the ubiquinol-cytochrome c reductase complex, 13 members of the cytochrome c oxidase complex, and 13 members of the ATP synthase complex), in 11 myosin light and heavy chain genes, in 8 genes of the mitochondrial ribosome complex, in 11 ribosomal protein-coding genes and in 12 genes encoding for solute carrier membrane transport proteins (**Files S4i-j**). Finally, of the top 100 most DEGs following retraining, 32 genes displayed transcriptional changes in the same direction as comparable RT studies of the metaMEx database and 76 genes in the opposite direction compared to metaMEx disuse studies (**File S4j**). Most genes within this top 100 were upregulated and related to metabolism (36), muscle structure and contraction (15) or muscle remodelling (10) (**File S4j**). This is the first study describing the retraining-related recovery of human muscle after disuse-induced atrophy in relation to the transcriptome and methylome. In rats, denervation-induced muscle atrophy decreased methylation in *MYOG, TRIM63, FBXO32* and *CHRNA1* together with increased expression that was returned to control levels following seven days of recovery.^63^ In accordance, *TRIM63, FBXO32* and *CHRNA1* were significantly upregulated following immobilization (FDR ≤ 0.067) in the present study and retraining reversed the RNA expression of all three genes to untrained and trained levels. We could not link this finding to any methylation changes.

Finally, pathway analysis on the methylome data identified 66, 79 and 12 enriched pathways (FDR < 0.1) following training, casting and retraining respectively (**Table 1, Files S4e-g**), of which 7 enriched during the whole intervention period. The overlapping pathways contained nearly the same number of hypo- and hypermethylated CpGs following training, whereas the frequency of hypermethylated CpGs clearly increased and decreased following casting and retraining, respectively (**Figure 2c**). The pathway analysis thereby confirmed that retraining induced more hypomethylation in comparison to training.

**Table 1.** Top 3 enriched pathways per analysis.

|  | ES | FDR | DMG | Genes in pathway | dmCpGs hypo | dmCpGs hyper |
| --- | --- | --- | --- | --- | --- | --- |
| **Young muscle following training\*** |  |  |  |  |  |  |
| - Longevity regulating | 11.1 | 0.005 | 27 | 89 | 9 | 30 |
| - Axon guidance | 9.7 | 0.009 | 43 | 182 | 24 | 23 |
| - Regulation of actin cytoskeleton | 9.4 | 0.009 | 48 | 213 | 35 | 23 |
| **Young muscle following casting\*** |  |  |  |  |  |  |
| - Focal adhesion | 17.8 | < 0.001 | 89 | 199 | 35 | 80 |
| - Prostate cancer | 13.0 | < 0.001 | 48 | 99 | 17 | 45 |
| - Rap1 signalling | 12.1 | < 0.001 | 84 | 206 | 38 | 69 |
| **Young muscle following retraining\*** |  |  |  |  |  |  |
| - MAPK signalling | 12.2 | 0.002 | 91 | 296 | 104 | 9 |
| - Focal adhesion | 8.5 | 0.031 | 61 | 199 | 61 | 12 |
| - Endocytosis | 8.1 | 0.032 | 72 | 247 | 75 | 16 |
| **Young versus older muscle** |  |  |  |  |  |  |
| - Focal adhesion | 37.4 | < 0.001 | 161 | 199 | 442 | 377 |
| - Axon guidance | 31.3 | < 0.001 | 145 | 182 | 356 | 447 |
| - Oxytocin signalling | 30.7 | < 0.001 | 125 | 153 | 372 | 314 |
| **Older muscle following training\*** |  |  |  |  |  |  |
| - Axon guidance | 12.9 | < 0.001 | 54 | 182 | 43 | 24 |
| - Cholinergic synapse | 11.0 | 0.003 | 36 | 112 | 27 | 20 |
| - Calcium signalling | 9.7 | 0.005 | 51 | 187 | 37 | 31 |
| **Older muscle following detraining\*** |  |  |  |  |  |  |
| - Focal adhesion | 19.6 | < 0.001 | 85 | 199 | 40 | 69 |
| - MAPK signalling | 10.7 | 0.004 | 102 | 296 | 41 | 88 |
| - Endometrial cancer | 9.6 | 0.004 | 28 | 59 | 13 | 23 |
| **Older muscle following retraining\*** |  |  |  |  |  |  |
| - Platelet activation | 14.3 | < 0.001 | 42 | 124 | 36 | 14 |
| - Phospholipase D signalling | 13.9 | < 0.001 | 47 | 147 | 40 | 16 |
| - Non-small cell lung cancer | 12.2 | < 0.001 | 26 | 67 | 21 | 11 |

ES = enrichment score; FDR = false discovery rate-adjusted p-value; DMG/dmCpGs = differentially methylated genes/CpG sites determined at the level of FDR < 0.1 for age-related analysis, p < 0.01 for other analyses. \* Compared to previous time point.

### Ageing-related hypermethylation in promoter-associated CpG islands and inverse expression levels

Comparing young to older muscle samples at baseline, we identified 50,828 age-related dmCpGs (FDR < 0.1, M-value difference > 0.4), of which 28,443 were hypomethylated and 22,385 were hypermethylated in older muscle tissue (**File S1a**). Two earlier genome-wide methylation studies^8,9^ that compared young and older human muscle samples found a clear global trend towards hypermethylated aged muscle (i.e. Turner et al., 2020 PREPRINT). We could only confirm this trend in promoter-associated CpG islands, where more than twice as many hypermethylated CpGs (2,761) were found compared to hypomethylated ones (1,202). In comparison, other regions contained 23 to 45 % more hypomethylated CpGs (**Figure 3a**). Promoter-associated dmCpGs in CpG islands were underrepresented compared to CpGs in the whole genome (**Figure 3b, File S1b**), which is a confirmation of the results of Zykovich et al.^8^, but in contrast to the findings of Turner et al.^9^, where most dmCpGs were located in CpG islands. Finally, we found a significant overlap of 2,334 CpGs between the age-related dmCpGs of the present study and the age-related dmCpGs of Zykovich et al. (p = 2.2e-16) (**Figure 3c, File S1d**). Interestingly, the age-related differences in β-values of the overlapping dmCpGs was nearly identical (r = 0.94, p < 0.001) (**Figure 3d**). These overlapping dmCpGs could, therefore, be a representation of an epigenetic clock^5^ common between individuals, whereas the dmCpGs uniquely found in each study represent the epigenetic drift^5^ between individuals due to stochastic factors and the specific environmental conditions of the population under investigation. To test this assumption, the 2,334 overlapping dmCpGs were compared with the 200 CpGs that were recently selected to create a muscle-specific epigenetic clock^64^. We found a significant overlap of 12 CpGs (p = 2.6e-12) (**Figure 3c, File S1d**).

**Figure 3.**
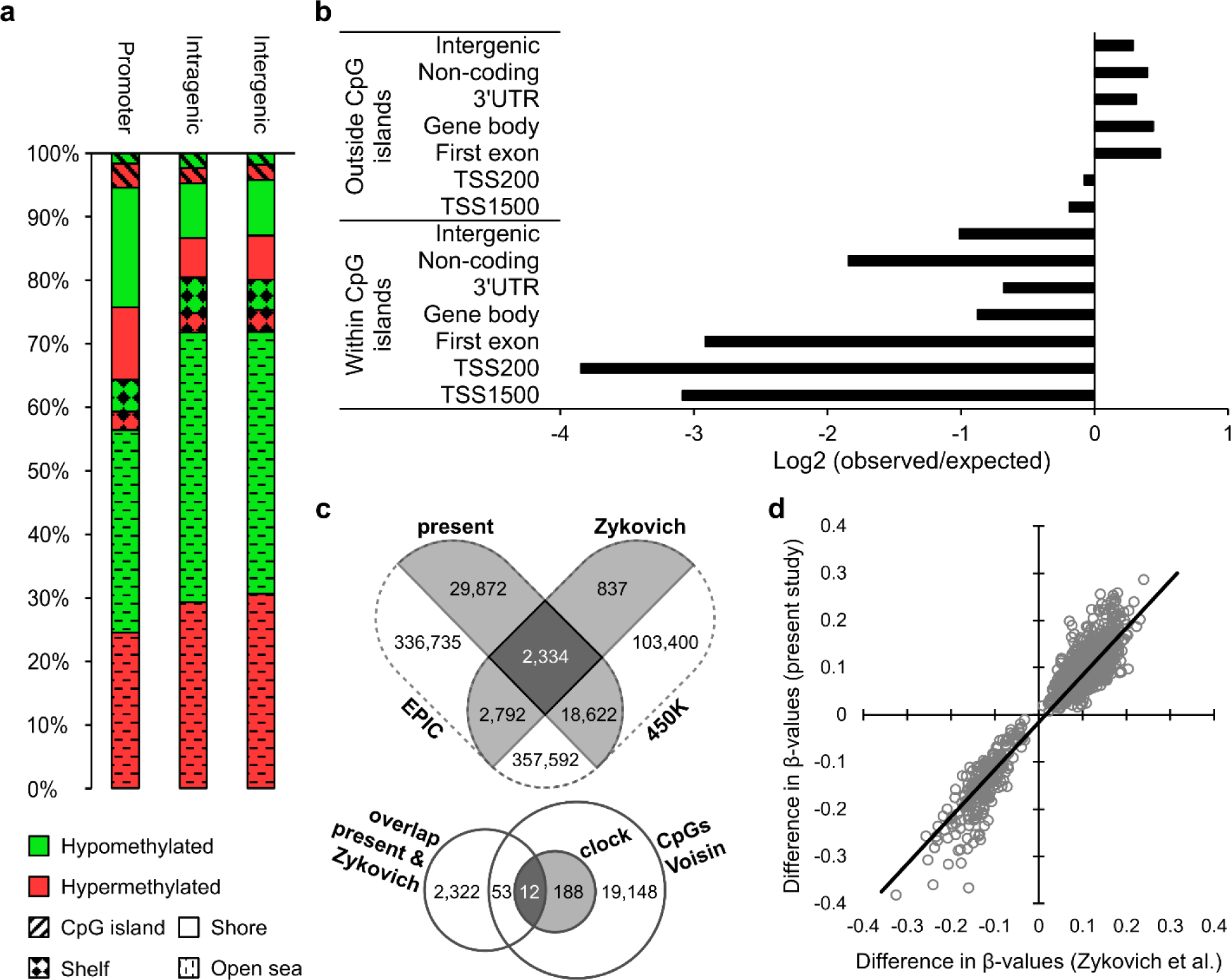
**(a)** Distribution of age-related dmCpGs (FDR < 0.1, M-value difference > 0.4) (older: n = 6, young: n = 5) across gene regions and structures, expressed as percentage of the total number per gene region. **(b)** Distribution of age-related dmCpGs across gene regions and structures (observed) relative to CpG distribution across the whole genome (expected). Negative log values correspond to an underrepresentation of dmCpGs within that area (positive values vice versa). **(c)** Top Venn-diagram displaying the overlap between the present age-related dmCpGs, selected from the MethylationEPIC Beadchip, and the age-related dmCpGs from Zykovich et al.^8^, selected from the HumanMethylation450K Beadchip. Bottom Venn-diagram displaying the overlap between the overlapping 2,334 dmCpGs from the top Venn-diagram and the 200 CpGs from the epigenetic clock of Voisin et al.^64^, selected from 19,401 CpGs. **(d)** Grey dots represent the difference between average β-values of older men and younger men per dmCpG (n = 2,334, cfr. c). The black line is the trend line.

Pathway analysis identified 151 enriched pathways (FDR < 0.1) (top 3 in **Table 1**), belonging to ‘organismal systems’ (52, e.g. axon guidance), ‘human diseases’ (44 of which 18 related to cancer), ‘environmental information processing’ (24, e.g. PI3K-Akt signalling pathway), ‘metabolism’ (16), ‘cellular processes’ (13, e.g. focal adhesion) and ‘genetic information processing’ (2) (**File S1c**).

In addition, ageing affected expression levels of 427 genes (FDR < 0.1), of which 183 were downregulated and 244 upregulated in older muscle. Remarkably, 71 % of these 427 DEGs contained at least one promoter-associated or intragenic dmCpG (FDR < 0.1, M-value difference > 0.4) (**File S1e**). We selected all DEGs with dmCpGs methylated in the same direction (i.e. all sites hypomethylated or all sites hypermethylated within a specific gene region) (**Figure 4a**, DEGs with differential methylation in promoter regions reported in **File S1f**). Most of the hypomethylated genes displayed increased expression, whereas expression of hypermethylated genes was generally downregulated. This was independent of the gene region and especially true for genes with more dmCpGs. However, many DEGs included both hypo- and hypermethylated CpGs (**File S1g**), which confirms that the relationship between methylation and expression is more nuanced than a simple inverse relation^65^. Exploring the top 100 differentially methylated and expressed genes between young and old, we found indications of impaired processes important for muscle functioning and remodelling in older muscle (**Table A2** in Appendix).

**Figure 4.**
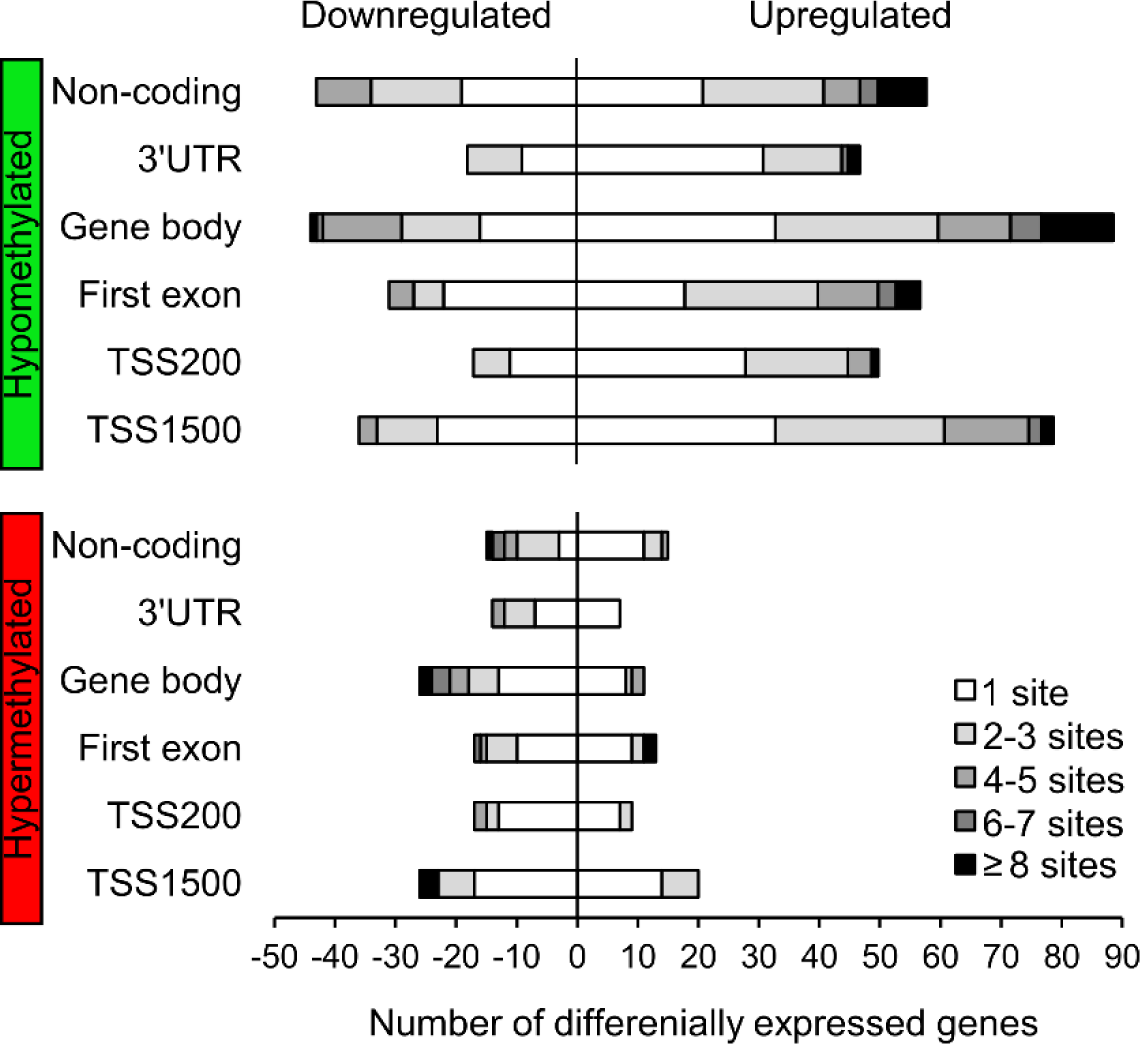
Number of hypomethylated (green) and hypermethylated (red) genes that displayed differential expression between young (n = 5) and older muscle (n = 6) (FDR < 0.1). Left of the Y-axes are downregulated genes, right are upregulated genes. Genes were categorized according to number of dmCpG sites, methylated in the same direction per gene region.

### Training has rejuvenating effects on the older muscle methylome

Twelve weeks of training in the older men rejuvenated their muscle methylome, as 37,339 of 50,828 age-related dmCpGs at baseline were no longer significantly different between trained older muscle and untrained young muscle (FDR ≥ 0.1 or M-value difference ≤ 0.4) (**File S1h**). Of these 37,346 CpGs, we selected the ones that changed significantly with training (606 CpGs with p < 0.01, M-value difference > 0.4). Based on these 606 CpGs, cluster analysis effectively clustered older post-training samples with younger baseline samples, instead of with older baseline samples (**Figure 5**). Pathway analysis linked four pathways to these CpGs (FDR < 0.1): ‘cortisol synthesis and secretion’ (ES = 9.6, FDR < 0.001), ‘cAMP signalling pathway’ (ES = 7.4, FDR = 0.067), ‘adrenergic signalling in cardiomyocytes’ (ES = 6.8, FDR = 0.067) and ‘aldosterone synthesis and secretion’ (ES = 6.75, FDR = 0.067). Most dmCpGs in these four pathways were hypermethylated in older compared to young muscle at baseline with a hyper/hypo ratio (i.e. number of hypermethylated CpGs/number of hypomethylated CpGs) between 2.7 and 11. Training had the opposite effect, directing the methylation level towards younger levels.

**Figure 5.**
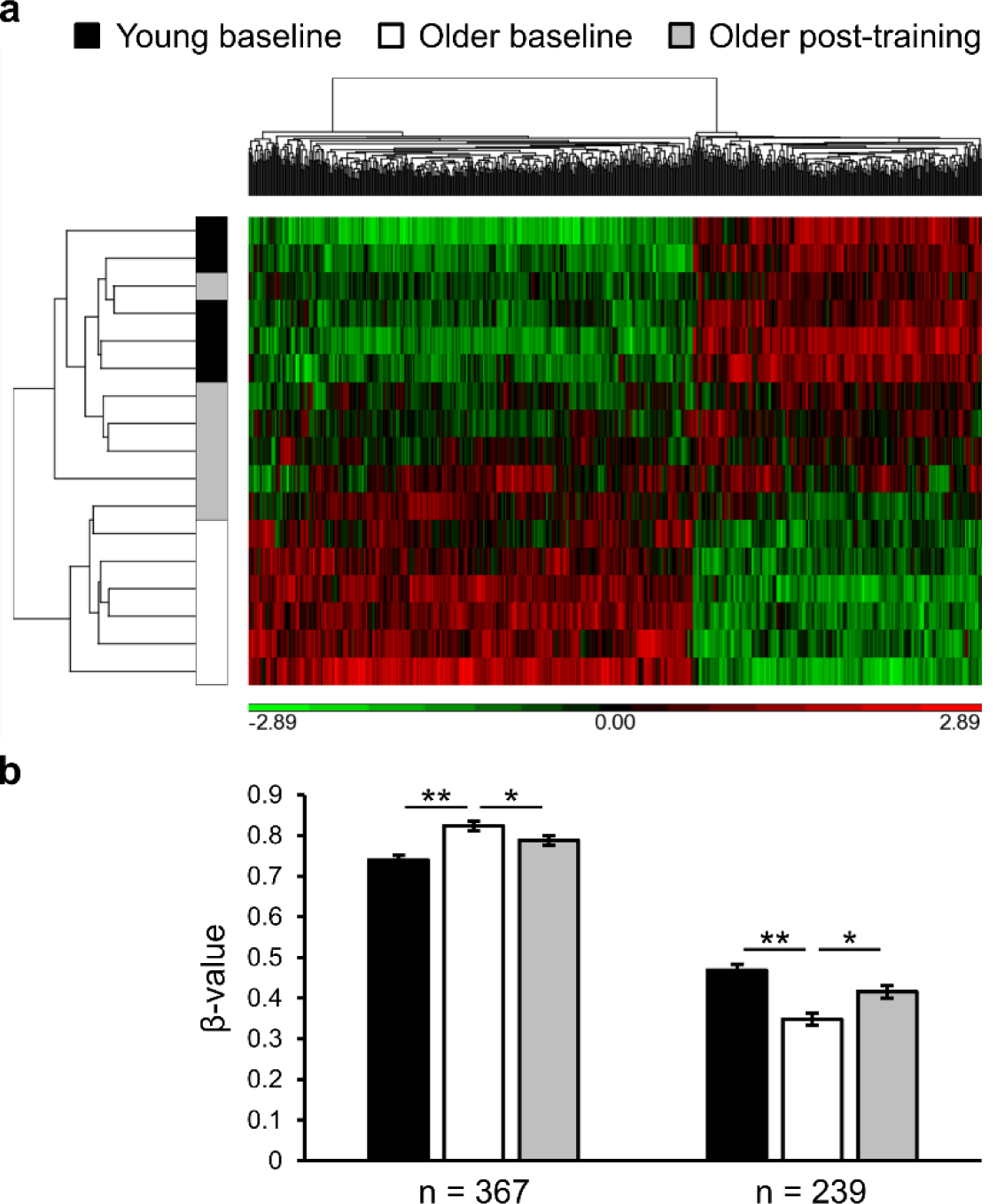
**(a)** Heat map of normalized β-values (shifted to mean of zero and scaled to SD of one) of CpGs (n = 606) that had significantly different methylation levels between young (n = 5, black) and older men at baseline (n = 6, white) and were reverted to younger levels following training in the older men (n = 6, grey). One CpG per column. Methylation levels are compared between subject groups (rows), with higher methylation coloured red and lower methylation coloured green. CpGs were hierarchically clustered in two categories: CpGs that are hypomethylated (left, n = 367) and CpGs that are hypermethylated (right, n = 239) following training in older men. **(b)** Average β-values of both clusters in each subject group. Values are mean ± SEM. ** FDR < 0.1 and difference between M-values > 0.4, * p < 0.01 and difference between M-values > 0.4.

In total, 2,657 CpGs were hypomethylated and 2,163 CpGs were hypermethylated after training compared to baseline (p < 0.01, M-value difference > 0.4) (**Figure 6a, File S2a**). In comparison, 12 weeks of normal daily living changed the methylation status of merely 255 CpGs in the older control group (p < 0.01, M-value difference > 0.4) (**Figure 6a, File S3a**). Especially in CpG islands and shores, training had a hypomethylating effect (**Figure 6b**). Comparable to the results in younger men, the dmCpGs were not over- or underrepresented in any region (**File S2d**). Of the 20 DEGs following training, only 1 displayed similar transcriptional changes following training as comparable training studies in older adults in the metaMEx meta-analysis (**File S2h**).

**Figure 6.**
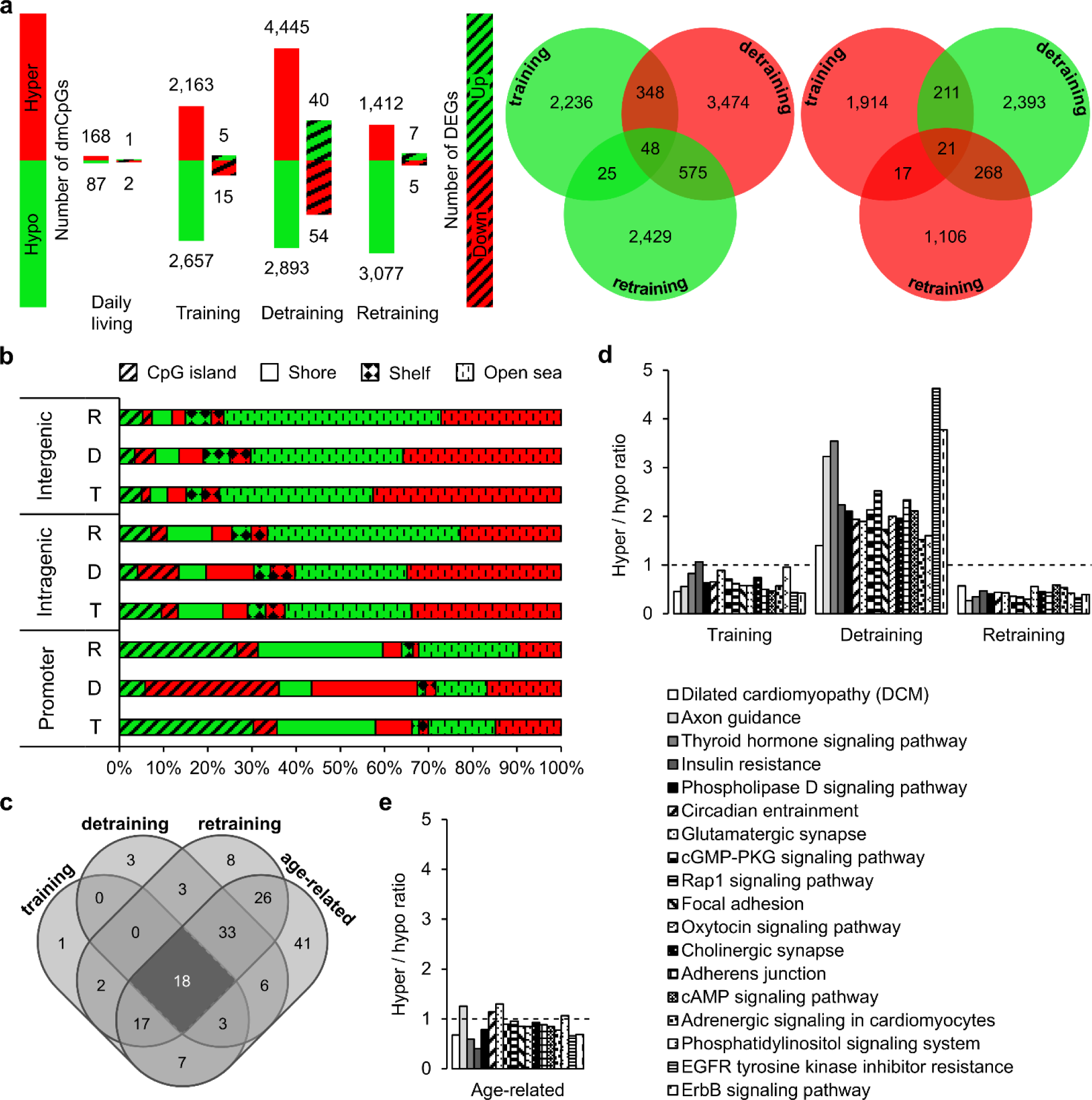
**(a)** Number of hypo- and hypermethylated CpGs (p < 0.01, M-value difference > 0.4; no pattern) versus up- and downregulated genes (DEGs) (FDR < 0.1, average read count ≥ 5; dashed pattern) following normal daily living (n = 3), training, detraining and retraining (n = 6/5, cfr. outlier RNA expression) compared to previous time point in older muscle. Venn-diagrams displaying overlap between dmCpGS following training, detraining and retraining (hypo- = green; hyper- = red) **(b)** Distribution of dmCpGs (% of the total, hypo- = green; hyper- = red) following training (T), detraining (D) and retraining (R) in older muscle across gene regions and gene structures. **(c)** Venn-diagram displaying the overlap of enriched pathways (FDR < 0.1) following training, detraining and retraining in older muscle and age-related pathways in older versus young muscle. **(d-e)** The hyper/hypo ratio of the 18 overlapping pathways identified in ‘c’. This ratio is calculated by dividing the number of hypermethylated CpGs with the number of hypomethylated CpGs. The dashed line (ratio = 1) delimitate an equal level of hyper- and hypomethylated CpGs within a pathway.

Against expectations, there were more genes downregulated than upregulated following a first training period (**Figure 6a**). Four of these genes (*PPDPFL, BBC3, PER1* and *MCL1*) were upregulated in older versus young muscle (FDR ≤ 0.068) and subsequently rejuvenated (i.e. downregulated) following training (FDR ≤ 0.038) (**Files S1e, S2h**). Twelve weeks of detraining, subsequently reverted the training effect and increased expression levels again in *PPDPFL, PER1* and *MCL1* (FDR ≤ 0.040) (**File S2i**). In addition, detraining reversed training adaptations in RNA expression of seven more genes (**File S2i**), as well as reverting methylation levels of 628 of the 4,820 dmCpGs following training (**Figure 6a, File S2b**). Globally, detraining mainly induced hypermethylation compared to post-training and more genes were downregulated (54) compared to upregulated (40) (**Figure 6a-b**). Eleven of these 94 DEGs were transcriptionally altered in the same direction as a five-day bed rest study in elderly men^56^, as displayed by the MetaMEx tool (**File S2i**). On the other hand, RNA expression of 10 genes changed in the opposite direction as this disuse study (**File S2i**), indicating that detraining and bed-rest might have a different effect on certain genes.

Similar to the results in young men, retraining hypomethylated more CpGs compared to training (3,077 vs. 2,657) (**Figure 6a**), which was observed across all gene regions and structures (**Figure 6b**). Pathway analysis identified 48, 66 and 107 enriched pathways (FDR < 0.1) following training, detraining and retraining, respectively (**Table 1, File S2e-g**). The majority of the pathways influenced by training, were also differentially methylated following detraining and/or retraining, whereas 34 pathways were uniquely affected by retraining (**Figure 6c**), suggesting that the methylome was more responsive to retraining. We found 18 pathways in common between each intervention period (**Figure 6c**). In the majority of the overlapping pathways, the number of hypomethylated CpGs further increased following retraining compared to training (**Figure 6d**). This could be indicative for an epi-memory within these pathways. Furthermore, these 18 pathways were also enriched when comparing the young and older muscle methylome at baseline (**Figure 6e**). Only 12 genes were transcriptionally altered following retraining (**Figure 6a, File S2j**). However, half of them were also picked up by comparable training studies of the MetaMEx meta-analysis and are involved in inflammation (*IL17D*), muscle growth inhibition (*PLN*), muscle contractions (*TPM3*), mechano-sensing (*ANKRD2*), muscle structure (*MYL5*) and energy metabolism (*CARNS1*) (FDR ≤ 0.091) (**File S2j**). Four other upregulated genes, not picked up by the meta-analysis, are involved in myogenic differentiation (*SOX11*), energy metabolism (*AK1, PGK1*) and muscle development and functioning (*PDLIM7*) (FDR ≤ 0.097) (**File S2j**).

### Genes indicative for an epi-memory in young and older skeletal muscle

Of the differentially expressed genes across all intervention phases in young and older muscle (FDR < 0.1, average read count ≥ 5), 11 to 33 % contained at least one dmCpG in the promoter or intragenic region (p < 0.01, M-value difference > 0.4) (**Figure 7**). Detailed information on these genes, including plots, is available in **Files S2l-m, S4l-m**. The interaction between the transcriptome and methylome was stronger following retraining compared to training. This might indicate that other epigenetic mechanisms were responsible for the alterations in chromatin accessibility during the first training period, whereas methylation was used during retraining to stabilize these changes.

**Figure 7.**
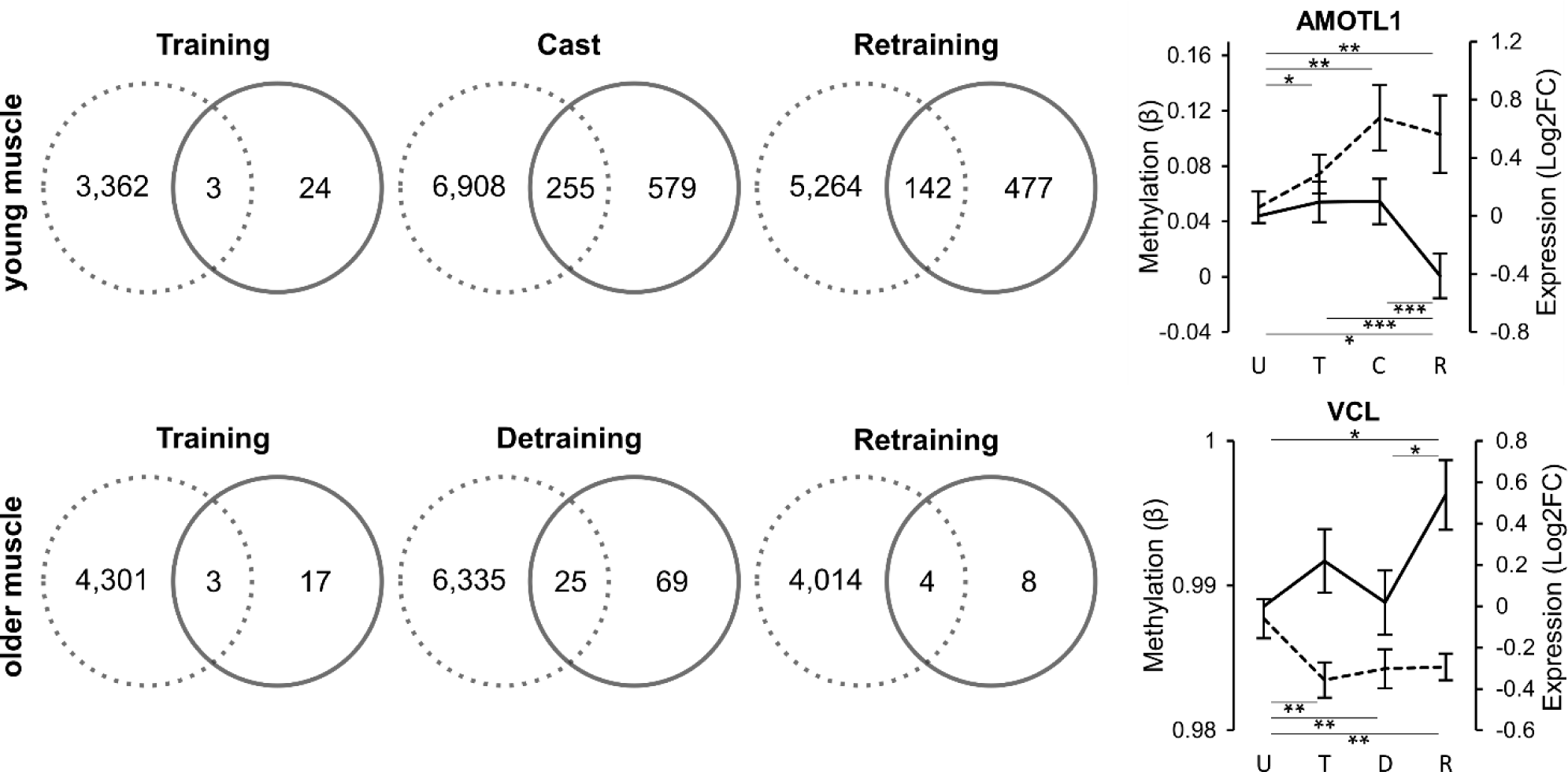
Overlap between differentially methylated (dotted) and expressed (full) genes following training (T), casting (C) / detraining (D) and retraining (R) compared to the previous time point in young (n=5/4, cfr. drop out after week 12) and older muscle (n = 6/5, cfr. outlier post-retraining RNA expression). *AMOTL1* and *VCL* are genes indicative for an epimemory in young and older muscle, respectively. Methylation values (dotted) are model-adjusted means ± SEM. Expression values (full) are model-adjusted log2 fold change values compared to baseline ± SEM. * p < 0.01, ** p < 0.001, *** FDR < 0.1.

We were specifically interested in genes constituting the epi-memory. Epi-memory genes can be defined as genes displaying an enhanced transcriptional response to retraining compared to training that could be linked to retained methylation modifications from the first training period (genes with retained methylation levels over time displayed in **Files S2k, S4k**). In young muscle, one CpG (cg15758225) within the open sea area of intron 3 of *AMOTL1* became more methylated following training (p = 0.006), which persisted during immobilization (p < 0.001) and retraining (p < 0.001) (**Figure 7**). As response to the second training period and potentially the hypermethylated state of cg15758225 pre-retraining, *AMOTL1* expression levels decreased (FDR = 0.012). This might indicate an epi-memory within *AMOTL1* and we found confirmation in the muscle biopsy data from Seaborne et al.^16^, as the hypermethylated state of two CpGs (cg13554187 and cg06447552) within *AMOTL1* following loading, was retained following unloading and for cg13554187 also following reloading. The investigators found several retraining-upregulated genes themselves indicative for an epi-memory in young muscle (*AXIN1, TRAF1, GRIK2, CAMK4* in Seaborn et al.^16^ and *FLNB, MYH9, SRGAP1, SRGN, ZMIZ1* in Turner et al.^27^). Despite finding one to three hypomethylated CpGs following retraining in *FLNB, MYH9, ZMIZ1, GRIK2* and *CAMK4*, we did not find the expected increased expression levels in these genes. We did find several genes with a trend towards enhanced expression levels following retraining and a correlated methylation pattern in young muscle (e.g. *NEXN, SERF2, PLCL2*) (**File S4m**).

In older men, we identified *VCL* as a gene with retained epigenetic modifications (**Figure 7**). Promoter- and intron-associated cg15758225, located in open sea area, was less methylated after training, detraining and retraining compared to baseline (all p < 0.001). This methylation change might have contributed to the trend towards an enhanced expression level following retraining (FDR = 0.136, p = 0.002). In addition, we found several other genes with a trend towards enhanced expression levels following retraining and a correlated methylation pattern in older muscle (e.g. *SOX11, PDLIM7, PGK1*) (**Files S2m**).

## DISCUSSION

In this study, we identified CpG methylation as a potential upstream mechanism that can explain transcriptional changes in skeletal muscle tissue associated with ageing and training. Interestingly, recurrent RT was accompanied with increasing numbers of hypomethylated CpGs and upregulated genes in young and older skeletal muscle and rejuvenated methylation levels of several CpGs altered by ageing. In addition, we found indications of an epi-memory within *AMOTL1* in young muscle and *VCL* in older muscle.

There have been many studies trying to reveal the age-associated human muscle transcriptome signature^66–69^, but literature lacks the integration of the mechanisms behind these transcriptional changes. We discovered that of the 427 DEGs between young and old, 71 % contained at least one dmCpG. This confirms, for the first time, that DNA methylation might be an important regulator of gene expression in muscle tissue. Furthermore, we found that the more dmCpGs methylated in the same direction, the clearer the inverse relationship between DNA methylation and RNA expression and this was regardless of the location of the dmCpG within the gene. It has been reported that a threshold of three CpGs is required to effectively inhibit transcription, with more CpGs and closer proximity to the promoter region associated with a higher percentage of gene repression.^70^ This explains why some but not all methylome adaptations over time mediated gene expression alterations in the present study, something that has been observed before in blood tissue^71,72^. In addition, the relationship between methylation and expression is more complex than a simple inverse correlation, as evidenced by the combination of both hypo- and hypermethylated CpGs within the different gene regions and structures of some genes. The complex transcriptome-methylome relationship has been described before^65^, but lacked tissue specificity. Therefore, we warrant further investigation into the interplay between methylation and other epigenetic mechanisms in muscle tissue (e.g. histone deacetylation, which works closely together with DNA methylation to silence genes^70^) and their role in the different gene regions and structures. Interestingly, older muscle tissue displayed signs of impaired myogenesis, mitochondrial bioenergetics, autophagy and glucose uptake, as well as indications of muscle atrophy and inflammageing at methylome and transcriptome level. These signs are closely related to cellular senescence^73^, one of the hallmarks of ageing^74^, and might contribute to age-related anabolic resistance and impaired skeletal muscle growth in response to an anabolic stimulus^75,76^. In accordance, we reported earlier that a first training period was unable to induce significant muscle growth in these same older men^28^. In the present study, we were able to link this impaired muscle growth to ageing-induced molecular dysregulation, both at transcriptional and epigenetic level.

Remarkably, 73 % of the age-related dmCpGs in untrained older muscle were no longer differentially methylated following 12 weeks of RT compared to young untrained muscle. The ability of resistance exercises to reverse age-related impairments have been extensively described at the muscle phenotypic level^77–79^, proteome level^52^, transcriptome level^54,60^ and other hallmarks of ageing^80^. For the first time, we confirm this finding at methylome level. We are aware of only one study^52^ that investigated the effect of RT on the older muscle methylome. However, the investigators were unable to find methylation changes after 12 weeks of RT, probably due to the low sample size, the strict FDR adjustment and the limited number of CpGs tested. We hypothesized that training would reverse the ageing-induced hypermethylation, predominantly found in older muscle tissue in literature^8,9^, based on the hypomethylating effect of aerobic exercise^10–13^, lifelong physical activity^15^ and RT in young to middle-aged adults^13,81^. However, both in young and older muscle tissue we observed that a first training period induced both hypomethylation and hypermethylation. That said, mainly training-induced hypomethylation was found in CpG islands and shores, possibly counteracting the ageing-induced hypermethylation in promoter-associated CpG islands of the older muscle samples. At transcriptional level, training altered 20 genes in older men. Interestingly, *PPDPFL, BBC3, PER1* and *MCL1* were upregulated in older versus young muscle, downregulated following training and upregulated again during detraining in older men. The role of *PPDPFL* in muscular training adaptations is currently unknown. The other three genes, on the other hand, are known to be involved in apoptosis (*BBC3*, BCL2 Binding Component 3, also known as *PUMA*)^82^ (*MCL1*, MCL1 Apoptosis Regulator, BCL2 Family Member)^83^ and circadian clock-regulated physiological processes (e.g. muscle hypertrophy) (*PER1*, Period Circadian Regulator 1)^84^.

Both young and older muscle responded better to a second training period. This was reflected by the higher number of hypomethylated CpGs and upregulated genes following retraining compared to training. In addition, retraining induced more hypomethylation than training did in the 7 overlapping pathways between training, immobilization and retraining in young men and in the 18 overlapping pathways between training, detraining and retraining in older men. Interestingly, these 18 pathways were also enriched in the age-related analysis, which supports the fact that chronic RT may act on several ageing-related pathways. Accordingly, the indications of inflammageing at baseline in older muscle tissue were only improved by the second training period, and not by the first. We observed nine hypomethylated dmCpGs in proximity to the promoter region of the pro-inflammatory cytokine Interleukin 17D (*IL17D*) when comparing older to young muscle, which was associated with elevated *IL17D* RNA expression in older muscle. *IL17D*, which also activates the acute inflammatory response mediated by NF-κB in the first hours after intense RT^85^, was successfully downregulated following retraining compared to post-detraining, whereas the decline in expression was not significant following the first training period. In addition, we found one hypermethylated dmCpG within *IL17D* following retraining. In agreement, NFKB Inhibitor Alpha (*NFKBIA*), inhibitor of NF-κB signalling, was downregulated following 12 weeks training. *NFKBIA* downregulation could be indicative for a permanent state of inflammation in older exercise-stimulated muscles. Surprisingly, *NFKBIA* expression levels returned to baseline levels during detraining and remained stable during retraining. These results indicate that previously trained older muscle may be less susceptible to exercise-induced chronic inflammation and recurrent RT may in fact improve age-related low grade inflammation, as has been reported before^86^.

Most interestingly, we identified two genes, indicative for an epi-memory within young and older muscle. In young men, the epi-memory pattern was reflected within the Angiomotin Like 1 (*AMOTL1*) gene. Upon activation of satellite cells, *AMOTL1* activates *YAP1* which triggers pro-proliferating genes and inhibits differentiation of myotubes.^87^ The decreased expression following retraining might signal myotube differentiation. Importantly, this result was confirmed when investigating the data of Seaborne et al.^16^. In older men, an increased expression level in Vinculin (*VCL*) following retraining was linked to the retained hypomethylated state of one CpG during training, detraining and retraining. In striated muscle, vinculin is a structural component of costameres^88^, which connects sarcomeres to the cell membrane to stabilize myofibres during contraction and relaxation. The gene, furthermore, potentiates mechanosensing^89,90^ and is upregulated in skeletal muscle tissue following chronic stimulation and disuse^91^. Most interestingly, *VCL* is part of the focal adhesion pathway, which was enriched during both training, detraining and retraining in older men, as well as during training, immobilization and retraining in younger men and the age-related comparison between young and old. Focal adhesion translates the (exercise-induced) cytoskeletal stress signals into cell growth by activating a cascade of signalling pathways.^92^ The importance of the focal adhesion pathway in our results might indicate that the rapid regain of training adaptations is related to improved mechano-sensing signalling during retraining. To support our results, we investigated other genes related to the mechanosensing pathway and found a trend towards enhanced gene expression and several hypomethylated CpGs following retraining in mechanosensitive transcriptional cofactor-encoding *WWTR1*, stretch-activated ion channel-encoding *PIEZO2* and extracellular-matrix associated *COL1A1* (**Figure A8** in Appendix).

Finally, we would like to address several limitations and future perspectives. First of all, sample sizes were small. This limited the statistical power and especially the intervention-related analyses might have been prone to type I error. Findings are consequently exploratory in nature and should be validated in future studies. On the other hand, findings that were found significant at FDR level of 0.1 despite the limited sample size, should be considered highly interesting. Secondly, CpGs may overlap with different regions of different gene transcripts belonging to one gene and it is not clear how each gene transcript is influenced by differential methylation, as RNA expression was only analysed at gene level. Thirdly, although we did perform SVA, we did not directly correct for cell type heterogeneity and fibre type differences between muscle samples. This might have confounded the analysis^93^. Finally, while we do integrate methylome and transcriptome data, we currently lack the translation to the proteome level, which should be the focus in future research. Also needed are similar analyses at single-cell or -nucleus level and the integration of other epigenetic mechanisms to unravel the precise role of epigenetics in the muscle memory phenomenon.

To conclude, these results indicate an enhanced transcriptional training response in previously trained young and older muscle, partially explained by retained and novel methylation adaptations. This supports that we might benefit from earlier training periods and that RT is an effective strategy to rejuvenate our methylome.

## SUPPORTING INFORMATION FILES

Supporting information files S1-4 are available on request.

## APPENDIX

**Table A2.**
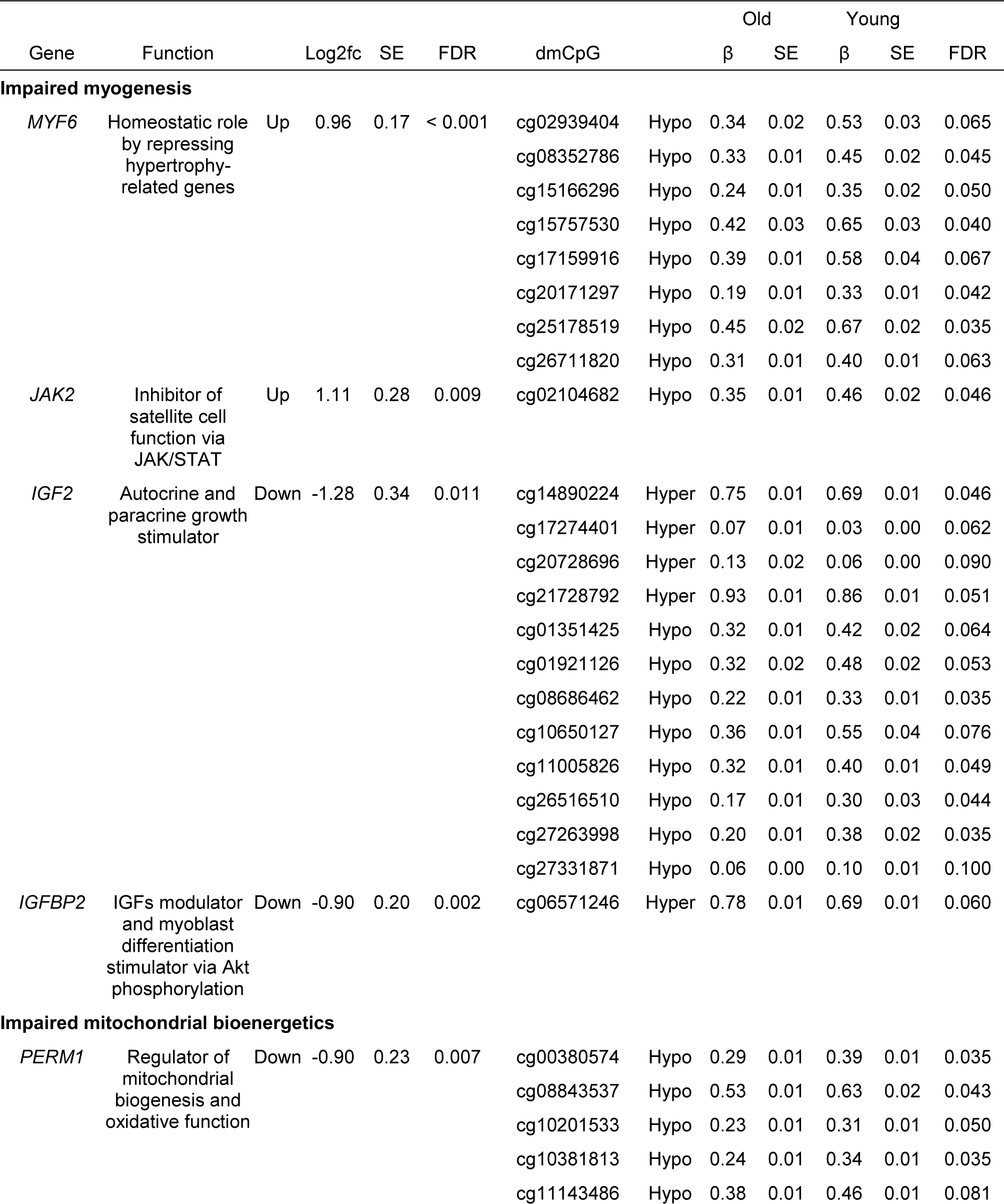
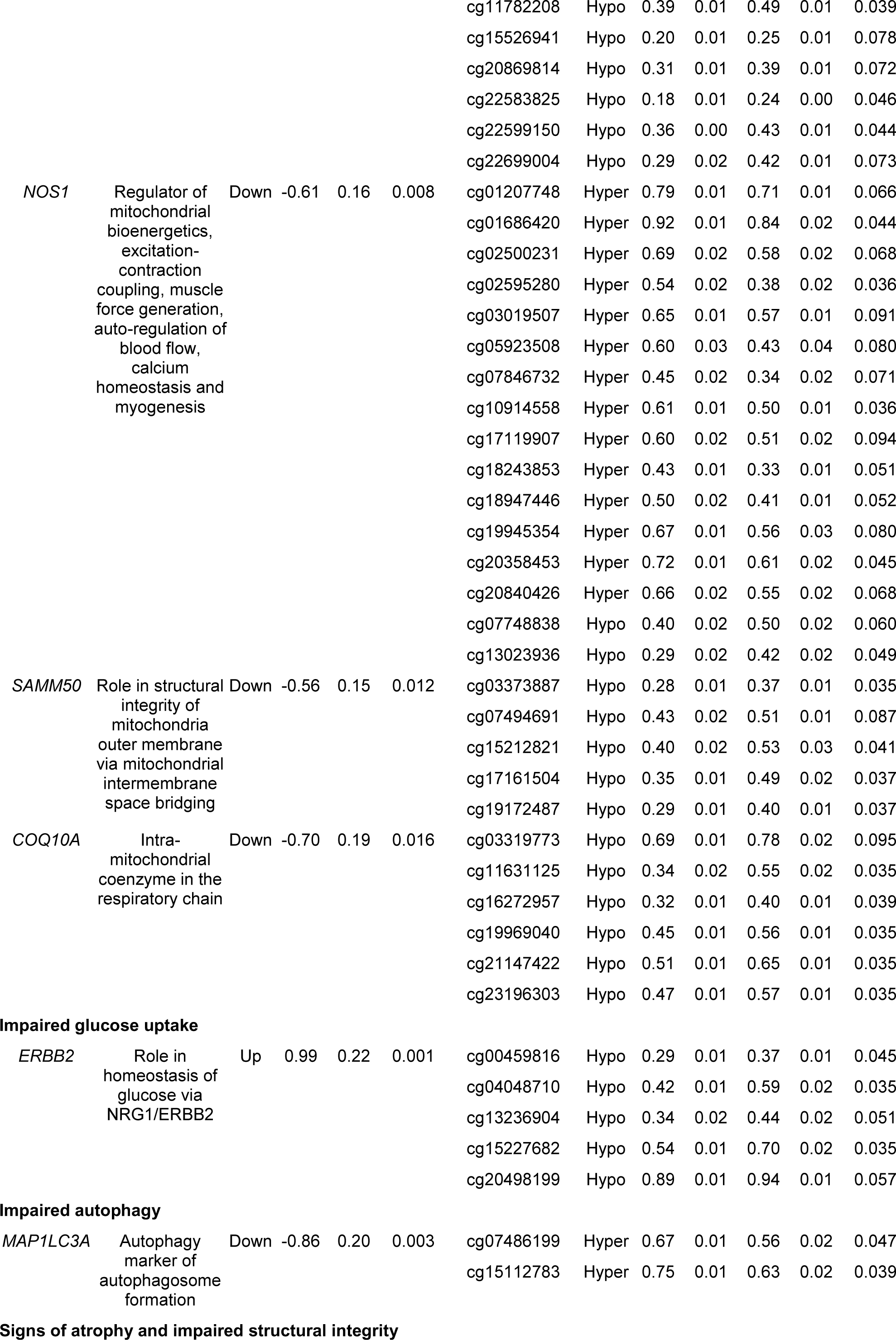
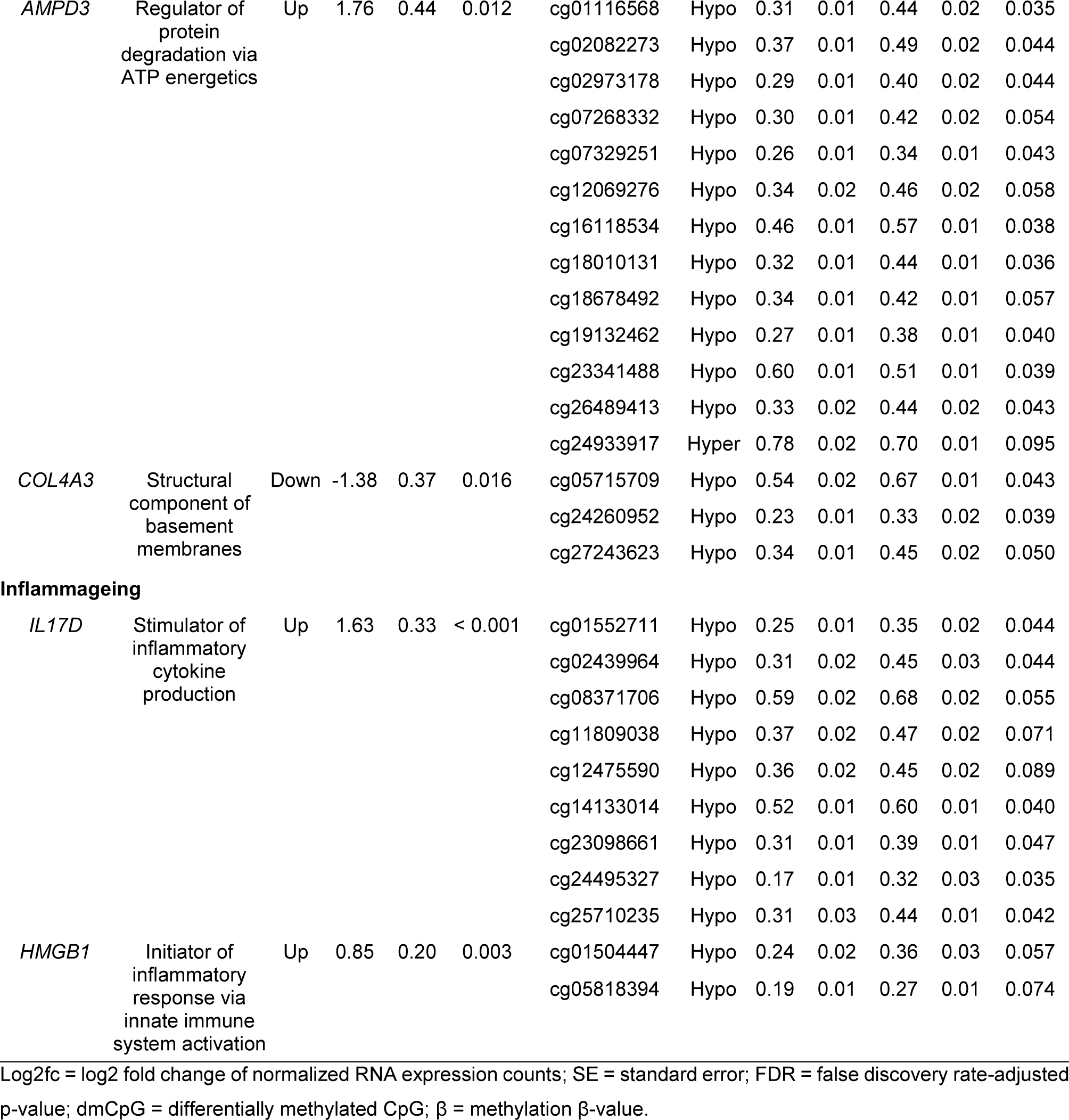
Important muscle-functioning genes differentially expressed and methylated in older muscle.

**Figure A8.**
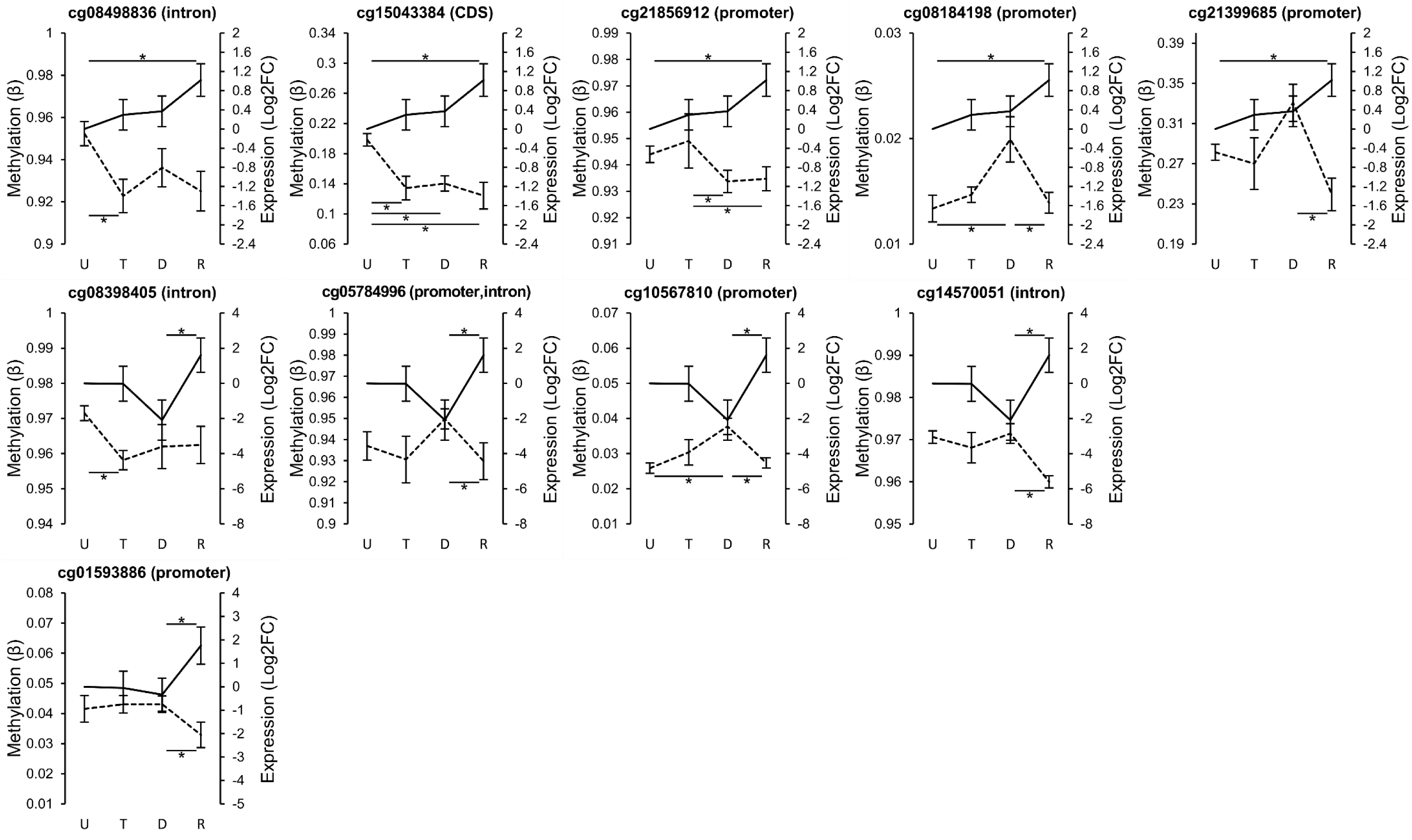
Enhanced expression following retraining and inversely correlated methylation levels in mechanosensing-related genes *WWTR1, PIEZO2* and *COL1A1*. Data from untrained (U) trained (T), detrained (D) and retrained (R) older muscle samples (n = 6/5, cfr. outlier post-retraining RNA expression). Methylation values (dotted) are model-adjusted means ± SEM. Expression values (full) are model-adjusted log2 fold change values compared to baseline ± SEM. * p < 0.01.

## ACKNOWLEDGMENTS

This study was supported by a research project grant (FWO G.0898.15) from Research Foundations Flanders. The authors thank Dr. Lingxiao He for sharing his DNA methylation processing script, Mr. Alvaro Cortes Calabuig for initial help with the processing of RNA expression data, Ms. Vanessa Brys for her help with the bioanalyzer.

Image acknowledgement: image of the muscle and DNA helix in Fig. 1 made by Freepik from www.flaticon.com.

## CONFLICT OF INTEREST

The authors report no conflict of interest.

## Notes

### Competing Interest Statement

The authors have declared no competing interest.

## BIBLIOGRAPHY

1. Cartee GD, Hepple RT, Bamman MM, Zierath JR. Exercise Promotes Healthy Aging of Skeletal Muscle. Cell Metab 2016;23:1034–1047.

2. Garatachea N, Lucía A. Genes and the ageing muscle: a review on genetic association studies. Age (Dordr) 2013;35:207–233.

3. Greenberg MVC, Bourc’his D. The diverse roles of DNA methylation in mammalian development and disease. Nat Rev Mol Cell Biol 2019;20:590–607.

4. Weber M, Hellmann I, Stadler MB, Ramos L, Pääbo S, Rebhan M et al. Distribution, silencing potential and evolutionary impact of promoter DNA methylation in the human genome. Nat Genet 2007;39:457–466.

5. Jones MJ, Goodman SJ, Kobor MS. DNA methylation and healthy human aging. Aging Cell 2015;14:924–932.

6. Pal S, Tyler JK. Epigenetics and aging. Sci Adv 2016;2:e1600584.

7. Day K, Waite LL, Thalacker-mercer A, West A, Bamman MM, Brooks JD et al. Differential DNA methylation with age displays both common and dynamic features across human tissues that are influenced by CpG landscape. Genome Biol 2013;14:R102.

8. Zykovich A, Hubbard A, Flynn JM, Tarnopolsky M, Fraga MF, Kerksick C et al. Genome-wide DNA methylation changes with age in disease-free human skeletal muscle. Aging Cell 2014;13:360–366.

9. Turner DC, Gorski PP, Maasar MF, Seaborne P, Baumert P, Brown AD et al. DNA methylation across the genome in aged human skeletal muscle tissue and muscle stem cells: The role of HOX genes and physical activity. bioRxiv [preprint], January 3, 2020 doi: 10.1101/2019.12.27.886135.

10. Alibegovic a C, Sonne MP, Højbjerre L, Jacobsen S, Nilsson E, Færch K et al. Insulin resistance induced by physical inactivity is associated with multiple transcriptional changes in skeletal muscle in young men. Am J Physiol Endocrinol Metab 2010;299:E752–E763.

11. Nitert MD, Dayeh T, Volkov P, Elgzyri T, Hall E, Nilsson E et al. Impact of an Exercise Intervention on DNA Methylation in Skeletal Muscle From First-Degree Relatives of Patients With Type 2 Diabetes. Diabetes 2012;61:3322–3332.

12. Lindholm ME, Marabita F, Gomez-Cabrero D, Rundqvist H, Ekström TJ, Tegnér J et al. An integrative analysis reveals coordinated reprogramming of the epigenome and the transcriptome in human skeletal muscle after training. Epigenetics 2014;9:37–41.

13. Rowlands DS, Page RA, Sukala WR, Giri M, Ghimbovschi SD, Hayat I et al. Multi-omic integrated networks connect DNA methylation and miRNA with skeletal muscle plasticity to chronic exercise in Type 2 diabetic obesity. Physiol Genomics 2014;46:747–765.

14. Laker RC, Lillard TS, Okutsu M, Zhang M, Hoehn KL, Connelly JJ et al. Exercise prevents maternal high-fat diet-induced hypermethylation of the Pgc-1a gene and age-dependent metabolic dysfunction in the offspring. Diabetes 2014;63:1605–1611.

15. Sailani MR, Halling JF, Møller HD, Lee H, Plomgaard P, Pilegaard H et al. Lifelong physical activity is associated with promoter hypomethylation of genes involved in metabolism, myogenesis, contractile properties and oxidative stress resistance in aged human skeletal muscle. Sci Rep 2019;9:3272.

16. Seaborne RA, Strauss J, Cocks M, Shepherd S, O’Brien TD, van Someren KA et al. Human skeletal muscle possesses an epigenetic memory of hypertrophy. Sci Rep 2018;8:1898.

17. Staron RS, Leonardi MJ, Karapondo DL, Malicky ES, Falkel JE, Hagerman FC et al. Strength and skeletal muscle adaptations in heavy-resistance-trained women after detraining and retraining. J Appl Physiol 1991;70:631–640.

18. Taaffe DR, Marcus R. Dynamic muscle strength alterations to detraining and retraining in elderly men. Clin Physiol 1997;17:311–324.

19. Henwood TR, Taaffe DR. Detraining and retraining in older adults Following Long-Term Muscle Power or Muscle Strength Specific Training. Journals Gerontol Ser A Biol Sci Med Sci 2008;63:751–758.

20. Moberg M, Lindholm ME, Reitzner SM, Ekblom B, Sundberg C-J, Psilander N. Exercise Induces Different Molecular Responses in Trained and Untrained Human Muscle. Med Sci Sport Exerc 2020 doi: 10.1249/mss.0000000000002310.

21. Gundersen K. Muscle memory and a new cellular model for muscle atrophy and hypertrophy. J Exp Biol 2016;219:235–242.

22. Jacobsen SC, Brøns C, Bork-Jensen J, Ribel-Madsen R, Yang B, Lara E et al. Effects of short-term high-fat overfeeding on genome-wide DNA methylation in the skeletal muscle of healthy young men. Diabetologia 2012;55:3341–3349.

23. Sharples AP, Stewart CE, Seaborne RA. Does skeletal muscle have an ‘epi’-memory? The role of epigenetics in nutritional programming, metabolic disease, aging and exercise. Aging Cell 2016;15:603–616.

24. Maples JM, Brault JJ, Shewchuk BM, Witczak CA, Zou K, Rowland N et al. Lipid exposure elicits differential responses in gene expression and DNA methylation in primary human skeletal muscle cells from severely obese women. Physiol Genomics 2015;47:139–146.

25. Sharples AP, Polydorou I, Hughes DC, Owens DJ, Hughes TM, Stewart CE. Skeletal muscle cells possess a ‘memory’ of acute early life TNF-a exposure : role of epigenetic adaptation. Biogerontology 2016;17:603–617.

26. Barrès R, Yan J, Egan B, Treebak JT, Rasmussen M, Fritz T et al. Acute Exercise Remodels Promoter Methylation in Human Skeletal Muscle. Cell Metab 2012;15:405–411.

27. Turner DC, Seaborne RA, Sharples AP. Comparative transcriptome and methylome analysis in human skeletal muscle anabolism, hypertrophy and epigenetic memory. Sci Rep 2019;9:4251.

28. Blocquiaux S, Gorski T, Roie E Van, Ramaekers M, Van Thienen R, Nielens H et al. The effect of resistance training, detraining and retraining on muscle strength and power, myofibre size, satellite cells and myonuclei in older men. Exp Gerontol 2020;133:110860.

29. Van Thienen R, Hulst GD, Deldicque L, Hespel P. Biochemical artifacts in experiments involving repeated biopsies in the same muscle. Physiol Rep 2014;2:e00286.

30. R Core Team. R: A language and environment for statistical computing. 2018 https://www.r-project.org/.

31. Aryee MJ, Jaffe AE, Corrada-Bravo H, Ladd-Acosta C, Feinberg AP, Hansen KD et al. Minfi: A flexible and comprehensive Bioconductor package for the analysis of Infinium DNA methylation microarrays. Bioinformatics 2014;30:1363–1369.

32. Wu MC, Kuan P-F. A Guide to Illumina BeadChip Data Analysis. In: Tost J, editor. Methods in Molecular Biology: DNA Methylation Protocols. Humana Press: New York; 2018. pp. 303–330.

33. Triche TJ, Weisenberger DJ, Van Den Berg D, Laird PW, Siegmund KD. Low-level processing of Illumina Infinium DNA Methylation BeadArrays. Nucleic Acids Res 2013;41:e90.

34. Bibikova M, Lin Z, Zhou L, Chudin E, Garcia EW, Wu B et al. High-throughput DNA methylation profiling using universal bead arrays. Genome Res 2006;16:383–393.

35. Teschendorff AE, Marabita F, Lechner M, Bartlett T, Tegner J, Gomez-Cabrero D et al. A beta-mixture quantile normalization method for correcting probe design bias in Illumina Infinium 450 k DNA methylation data. Bioinformatics 2013;29:189–196.

36. Pidsley R, Y Wong CC, Volta M, Lunnon K, Mill J, Schalkwyk LC. A data-driven approach to preprocessing Illumina 450K methylation array data. BMC Genomics 2013;14:293.

37. Liu J, Siegmund KD. An evaluation of processing methods for HumanMethylation450 BeadChip data. BMC Genomics 2016;17:469.

38. Zhou W, Laird PW, Shen H. Comprehensive characterization, annotation and innovative use of Infinium DNA methylation BeadChip probes. Nucleic Acids Res 2017;45:e22.

39. Du P, Zhang X, Huang C-C, Jafari N, Kibbe WA, Hou L et al. Comparison of Beta-value and M-value methods for quantifying methylation levels by microarray analysis. BMC Bioinformatics 2010;11:587.

40. Leek JT, Johnson WE, Parker HS, Jaffe AE, Storey JD. The SVA package for removing batch effects and other unwanted variation in high-throughput experiments. Bioinformatics 2012;28:882–883.

41. Leek JT, Storey JD. Capturing heterogeneity in gene expression studies by surrogate variable analysis. PLoS Genet 2007;3:1724–1735.

42. Benjamini Y, Hochberg Y. Controlling the False Discovery Rate: A Practical and Powerful Approach to Multiple Testing. J R Stat Soc Ser B 1995;57:289–300.

43. Gaus W, Mayer B, Muche R. Interpretation of Statistical Significance - Exploratory Versus Confirmative Testing in Clinical Trials, Epidemiological Studies, Meta-Analyses and Toxicological Screening (Using Ginkgo biloba as an Example). J Clin Exp Pharmacol 2015;5:1000182.

44. Reimand J, Isser R, Voisin V, Kucera M, Tannus-lopes C, Rostamianfar A et al. Pathway enrichment analysis and visualization of omics data using g:Profiler, GSEA, Cytoscape and EnrichmentMap. Nat Protoc 2019;14:482–517.

45. Li H, Handsaker B, Wysoker A, Fennell T, Ruan J, Homer N et al. The Sequence Alignment/Map format and SAMtools. Bioinformatics 2009;25:2078–2079.

46. Anders S, Pyl PT, Huber W. HTSeq-A Python framework to work with high-throughput sequencing data. Bioinformatics 2015;31:166–169.

47. Love MI, Huber W, Anders S. Moderated estimation of fold change and dispersion for RNA-seq data with DESeq2. Genome Biol 2014;15:550.

48. Ignatiadis N, Klaus B, Zaugg JB, Huber W. Data-driven hypothesis weighting increases detection power in genome-scale multiple testing. Nat Methods 2016;13:577–580.

49. Pillon NJ, Gabriel BM, Dollet L, Smith JAB, Sardón Puig L, Botella J et al. Transcriptomic profiling of skeletal muscle adaptations to exercise and inactivity. Nat Commun 2020;11:470.

50. Phillips BE, Williams JP, Gustafsson T, Bouchard C, Rankinen T, Knudsen S et al. Molecular Networks of Human Muscle Adaptation to Exercise and Age. PLoS Genet 2013;9:e1003389.

51. Damas F, Ugrinowitsch C, Libardi CA, Jannig PR, Hector AJ, McGlory C et al. Resistance training in young men induces muscle transcriptome-wide changes associated with muscle structure and metabolism refining the response to exercise-induced stress. Eur J Appl Physiol 2018;118:2607–2616.

52. Robinson MM, Dasari S, Konopka AR, Johnson ML, Manjunatha S, Esponda RR et al. Enhanced Protein Translation Underlies Improved Metabolic and Physical Adaptations to Different Exercise Training Modes in Young and Old Humans. Cell Metab 2017;25:581–592.

53. Liu D, Sartor MA, Nader GA, Gutmann L, Treutelaar MK, Pistilli EE et al. Skeletal muscle gene expression in response to resistance exercise: sex specific regulation. BMC Genomics 2010;11:659.

54. Raue U, Trappe T a., Estrem ST, Qian H-R, Helvering LM, Smith RC et al. Transcriptome signature of resistance exercise adaptations: mixed muscle and fiber type specific profiles in young and old adults. J Appl Physiol 2012;112:1625–1636.

55. Gordon PM, Liu D, Sartor MA, Iglayreger HB, Pistilli EE, Gutmann L et al. Resistance exercise training influences skeletal muscle immune activation: a microarray analysis. J Appl Physiol 2012;112:443–453.

56. Mahmassani ZS, Reidy PT, McKenzie AI, Stubben C, Howard MT, Drummond MJ. Disuse-induced insulin resistance susceptibility coincides with a dysregulated skeletal muscle metabolic transcriptome. J Appl Physiol 2019;126:1419–1429.

57. Rullman E, Mekjavic IB, Fischer H, Eiken O. PlanHab (Planetary Habitat Simulation): the combined and separate effects of 21 days bed rest and hypoxic confinement on human skeletal muscle miRNA expression. Physiol Rep 2016;4:e12753.

58. Lammers G, Duijnhoven NTL Van, Hoenderop JG, Horstman AM, Haan A De, Janssen TWJ et al. The identification of genetic pathways involved in vascular adaptations after physical deconditioning versus exercise training in humans. Exp Physiol 2013;98:710–721.

59. Abadi A, Glover EI, Isfort RJ, Raha S, Safdar A, Yasuda N et al. Limb immobilization induces a coordinate down-regulation of mitochondrial and other metabolic pathways in men and women. PLoS One 2009;4:e6518.

60. Melov S, Tarnopolsky MA, Beckman K, Felkey K, Hubbard A. Resistance Exercise Reverses Aging in Human Skeletal Muscle. PLoS One 2007;2:e465.

61. Hangelbroek RWJ, Fazelzadeh P, Tieland M, Boekschoten M V., Hooiveld GJEJ, van Duynhoven JPM et al. Expression of protocadherin gamma in skeletal muscle tissue is associated with age and muscle weakness. J Cachexia Sarcopenia Muscle 2016;7:604–614.

62. Tarnopolsky M, Phillips S, Parise G, Varbanov A, DeMuth J, Stevens P et al. Gene expression, fiber type, and strength are similar between left and right legs in older adults. Journals Gerontol - Ser A Biol Sci Med Sci 2007;62:1088–1095.

63. Fisher AG, Seaborne RA, Hughes TM, Gutteridge A, Stewart C, Coulson JM et al. Transcriptomic and epigenetic regulation of disuse atrophy and the return to activity in skeletal muscle. FASEB J 2017;31:5268–5282.

64. Voisin S, Harvey NR, Haupt LM, Griffiths LR, Ashton KJ, Coffey VG et al. An epigenetic clock for human skeletal muscle. J Cachexia Sarcopenia Muscle 2020 doi: 10.1002/jcsm.12556.

65. Jones PA. Functions of DNA methylation: Islands, start sites, gene bodies and beyond. Nat Rev Genet 2012;13:484–492.

66. Zahn JM, Sonu R, Vogel H, Crane E, Mazan-Mamczarz K, Rabkin R et al. Transcriptional profiling of aging in human muscle reveals a common aging signature. PLoS Genet 2006;2:1058–1069.

67. Giresi PG, Stevenson EJ, Theilhaber J, Koncarevic A, Parkington J, Fielding RA et al. Identification of a molecular signature of sarcopenia. Physiol Genomics 2005;21:253–263.

68. Su J, Ekman C, Oskolkov N, Lahti L, Ström K, Brazma A et al. A novel atlas of gene expression in human skeletal muscle reveals molecular changes associated with aging. Skelet Muscle 2015;5:35.

69. Shafiee G, Asgari Y, Soltani A, Larijani B, Heshmat R. Identification of candidate genes and proteins in aging skeletal muscle (sarcopenia) using gene expression and structural analysis. PeerJ 2018;6:e5239.

70. Curradi M, Izzo A, Badaracco G, Landsberger N. Molecular Mechanisms of Gene Silencing Mediated by DNA Methylation. Mol Cell Biol 2002;22:3157–3173.

71. Yuan T, Jiao Y, de Jong S, Ophoff RA, Beck S, Teschendorff AE. An Integrative Multi-scale Analysis of the Dynamic DNA Methylation Landscape in Aging. PLoS Genet 2015;11:1–21.

72. Steegenga WT, Boekschoten M V., Lute C, Hooiveld GJ, De Groot PJ, Morris TJ et al. Genome-wide age-related changes in DNA methylation and gene expression in human PBMCs. Age (Omaha) 2014;36:1523–1540.

73. Ferrucci L, Fabbri E. Inflammageing: chronic inflammation in ageing, cardiovascular disease, and frailty. Nat Rev Cardiol 2018;15:505–522.

74. López-otín C, Blasco MA, Partridge L, Serrano M, Kroemer G. The Hallmarks of Aging Longevity. Cell 2013;153:1194–1217.

75. Mero AA, Hulmi JJ, Salmijärvi H, Katajavuori M, Haverinen M, Holviala J et al. Resistance training induced increase in muscle fiber size in young and older men. Eur J Appl Physiol 2013;113:641–650.

76. Tieland M, Trouwborst I, Clark BC. Skeletal muscle performance and ageing. J Cachexia Sarcopenia Muscle 2018;9:3–19.

77. Fragala MS, Cadore EL, Dorgo S, Izquierdo M, Kraemer WJ, Peterson MD et al. Resistance Training for Older Adults: Position Statement From the National Strength and Conditioning Association. J strength Cond Res 2019;33:2019–2052.

78. Candow D, Chilibeck P, Abeysekara S, Zello G. Short-term heavy resistance training eliminates age-related deficits in muscle mass and strength in healthy older males. J Strength Cond Res 2011;25:326–333.

79. McLeod M, Breen L, Hamilton DL, Philp A. Live strong and prosper: the importance of skeletal muscle strength for healthy ageing. Biogerontology 2016;17:497–510.

80. Rebelo-Marques A, Lages ADS, Andrade R, Ribeiro CF, Mota-Pinto A, Carrilho F et al. Aging hallmarks: The benefits of physical exercise. Front Endocrinol (Lausanne) 2018;9:258.

81. Lund J, Rustan AC, Løvsletten NG, Mudry JM, Langleite TM, Feng YZ et al. Exercise in vivo marks human myotubes in vitro: Training-induced increase in lipid metabolism. PLoS One 2017;12:e0175441.

82. Harford TJ, Kliment G, Shukla GC, Weyman CM. The muscle regulatory transcription factor MyoD participates with p53 to directly increase the expression of the pro-apoptotic Bcl2 family member PUMA. Apoptosis 2017;22:1532–1542.

83. Yang T, Buchan HL, Townsend KJ, Craig RW. MCL-1, a member of the BCL-2 family, is induced rapidly in response to signals for cell differentiation or death, but not to signals for cell proliferation. J Cell Physiol 1996;166:523–536.

84. Popov D V, Makhnovskii PA, Kurochkina NS, Lysenko EA, Vepkhvadze TF, Vinogradova OL. Intensity-dependent gene expression after aerobic exercise in endurance-trained skeletal muscle. Biol Sport 2018;35:277–289.

85. Vella L, Caldow MK, Larsen AE, Tassoni D, Gatta PAD, Gran P et al. Resistance exercise increases NF-κB activity in human skeletal muscle. Am J Physiol - Regul Integr Comp Physiol 2012;302:667–673.

86. Calle MC, Fernandez ML. Effects of resistance training on the inflammatory response. Nutr Res Pract 2010;4:259–269.

87. Li L, Fan C. A CREB-MPP7-AMOT regulatory axis controls muscle stem cell expansion and self-renewal competence. Cell Rep 2017;21:1253–1266.

88. Pardo J V., D’Angelo Siliciano J, Craig SW. A vinculin-containing cortical lattice in skeletal muscle: Transverse lattice elements (‘costameres’) mark sites of attachment between myofibrils and sarcolemma. Proc Natl Acad Sci U S A 1983;80:1008–1012.

89. Rindom E, Vissing K. Mechanosensitive molecular networks involved in transducing resistance exercise-signals into muscle protein accretion. Front Physiol 2016;7:547.

90. Chorev DS, Volberg T, Livne A, Eisenstein M, Martins B, Kam Z et al. Conformational states during vinculin unlocking differentially regulate focal adhesion properties. Sci Rep 2018;8:2693.

91. Rezvani M, Ornatsky O, Connor M, Eisenberg H, Hood D. Dystrophin, vinculin, and aciculin in skeletal muscle subject to chronic use and disuse. Med Sci Sport Exerc 1996;28:79–84.

92. Graham ZA, Gallagher PM, Cardozo CP. Focal adhesion kinase and its role in skeletal muscle. J Muscle Res Cell Motil 2015;36:305–315.

93. Taylor DL, Jackson AU, Narisu N, Hemani G, Erdos MR, Chines PS et al. Integrative analysis of gene expression, DNA methylation, physiological traits, and genetic variation in human skeletal muscle. Proc Natl Acad Sci U S A 2019;166:10883–10888.

