## Supporting information figures S1-3 for "Recurrent training rejuvenates and enhances transcriptome and methylome responses in young and older human muscle"

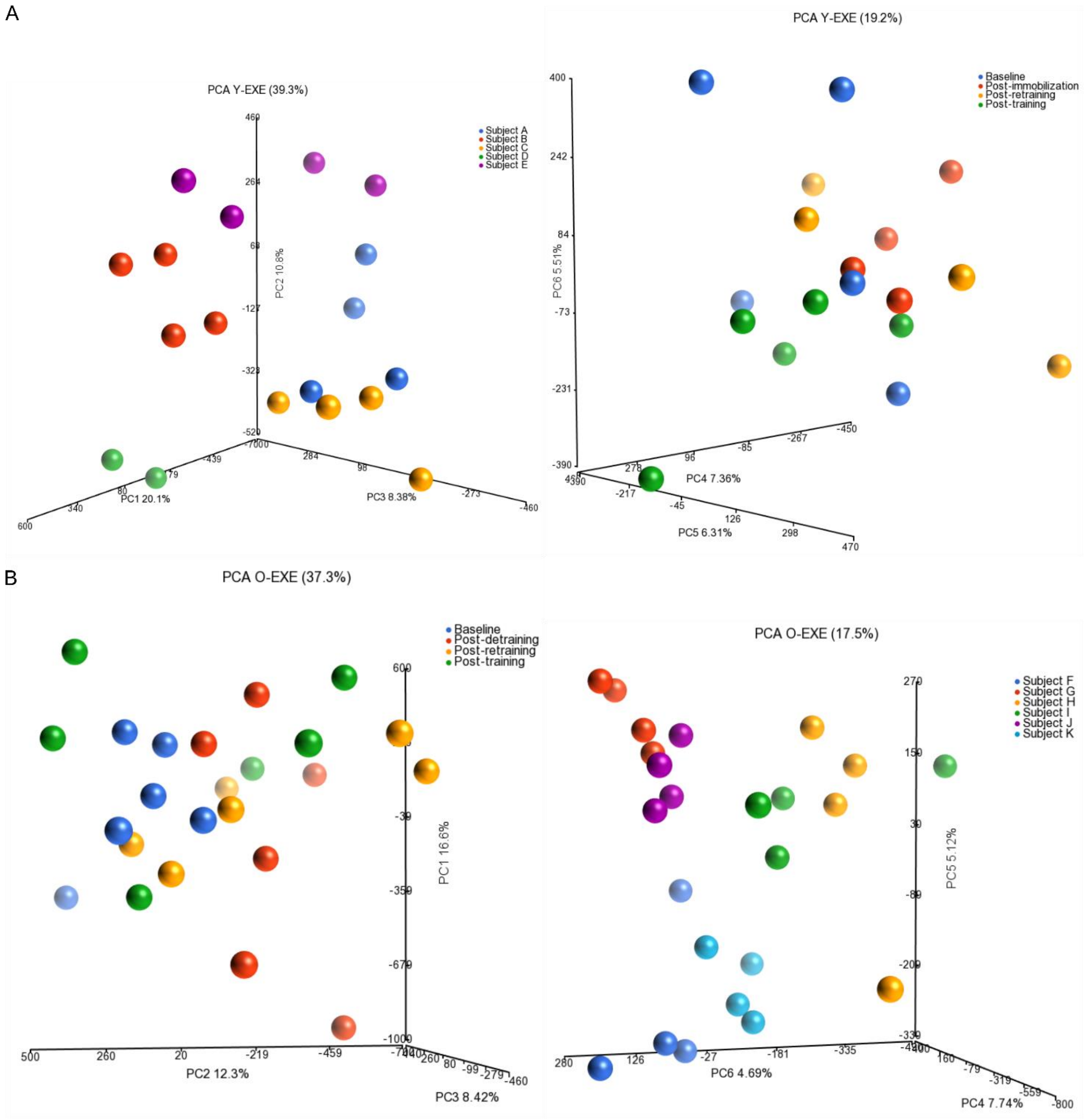


**Figure S1 Principal component analysis plots of methylation data in the intervention analysis.** Plots are depicted post-normalization and post-filtering. **(A)** Plots of young muscles. PC1-3 grouped samples based on sample ID. PC4-6 grouped samples based on timing in the intervention. **(B)** Plots of older muscles. PC1-3 clustered samples based on timing in the intervention. PC4-6 clustered samples based on sample ID.

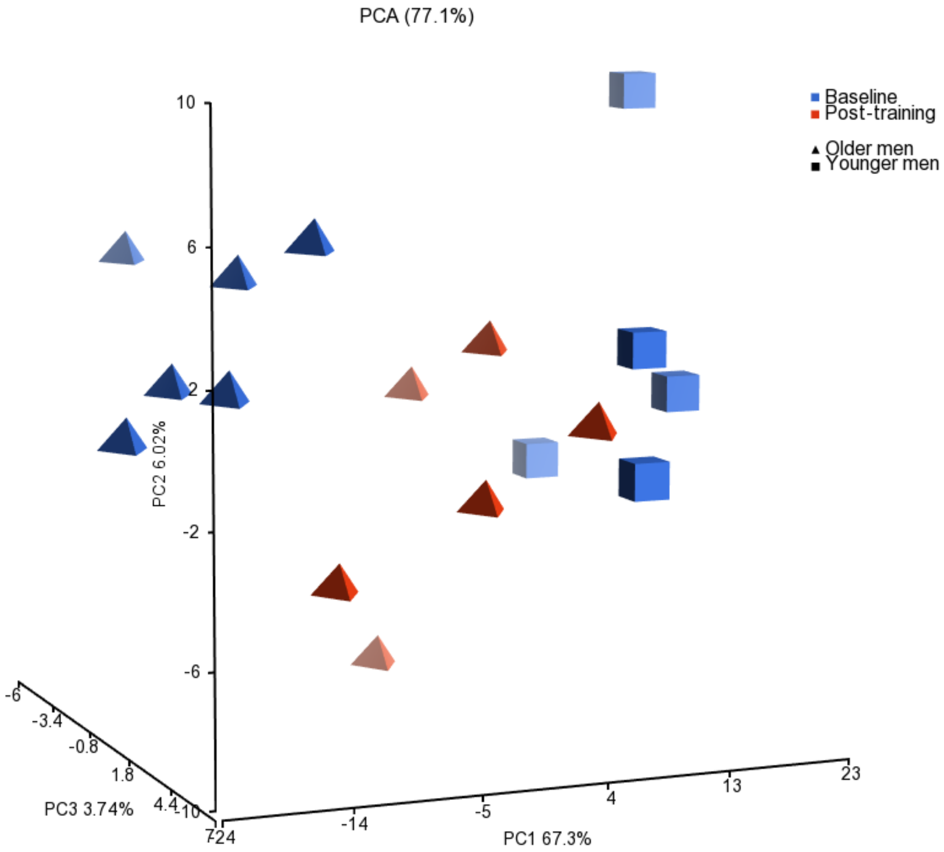


**Figure S2 Principal components analysis plots of methylation data in the age-related analysis.** The plot is depicted post-normalization and post-filtering. PCA1-3 clustered samples based on age (old versus young) and timing (baseline versus post-training)

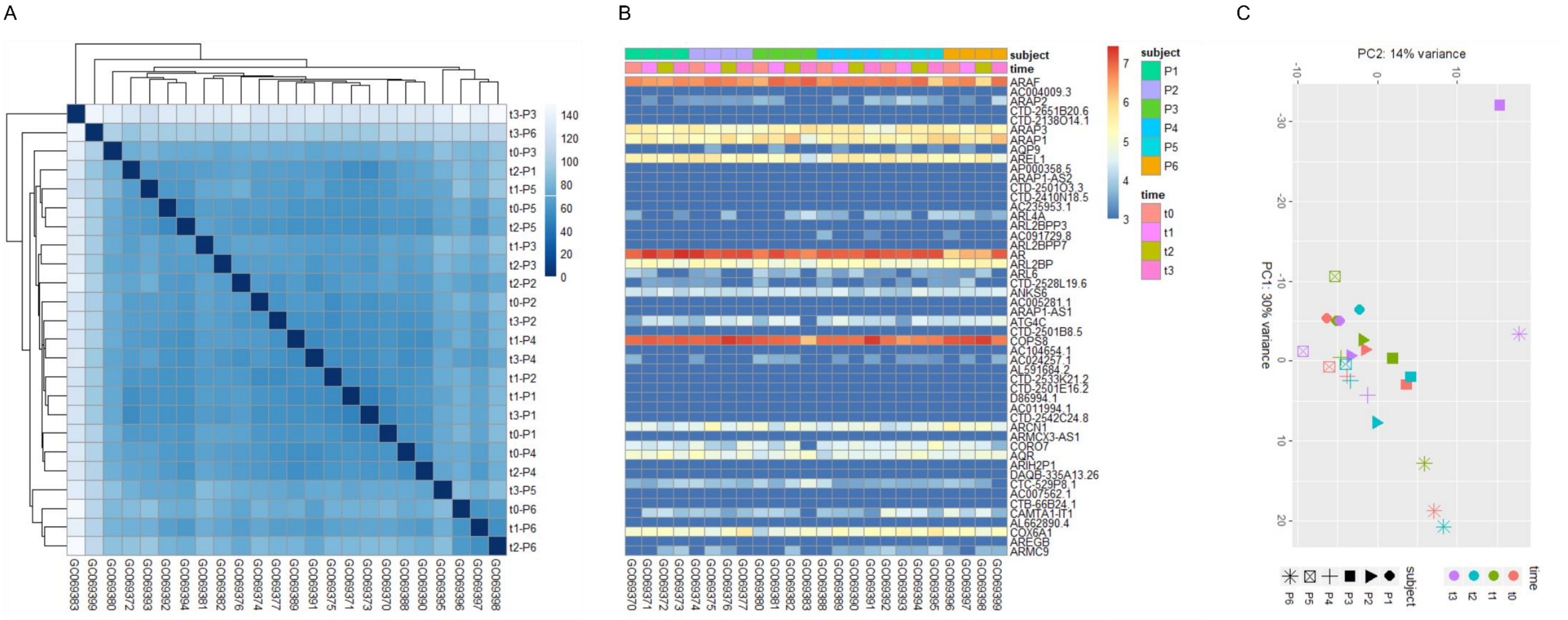


**Figure S3 Outlier detection in RNA expression data. (A)** Sample distance plot, **(B)** heatmap and **(C)** principal component plot all identified subject P3-t3 (GC069383) of the older exercise group as outlier. In addition, the total number of read counts of this sample was particularly low (< 500,000) compared to the other samples (> 1,000,000). Therefore, we decided to exclude this sample from the expression analysis.
